# Light-driven formation of high-valent manganese oxide by photosystem II supports evolutionary role in early bioenergetics

**DOI:** 10.1101/2020.03.03.975516

**Authors:** Petko Chernev, Sophie Fischer, Jutta Hoffmann, Nicholas Oliver, Robert L. Burnap, Ivelina Zaharieva, Dennis J. Nürnberg, Michael Haumann, Holger Dau

**Affiliations:** Physics Department, Freie Universität Berlin, Arnimallee 14, 14195 Berlin, Germany; Department of Chemistry - Ångström Laboratory, Molecular Biomimetics, Uppsala University, Lägerhyddsvägen 1, 75120 Uppsala, Sweden; Dept. of Microbiology and Molecular Genetics, Oklahoma State University, Stillwater, Oklahoma 74078-4034, United States

## Abstract

Water oxidation and concomitant O_2_-formation by the Mn_4_Ca cluster of oxygenic photosynthesis has shaped the biosphere, atmosphere, and geosphere. It has been hypothesized that at an early stage of evolution, before photosynthetic water oxidation became prominent, photosynthetic formation of Mn oxides from dissolved Mn(2+) ions may have played a key role in bioenergetics and possibly facilitated early geological manganese deposits. The biochemical evidence for the ability of photosystems to form extended Mn oxide particles, lacking until now, is provided herein. We tracked the light-driven redox processes in spinach photosystem II (PSII) particles devoid of the Mn_4_Ca clusters by UV-vis and X-ray spectroscopy. We find that oxidation of aqueous Mn(2+) ions results in PSII-bound Mn(III,IV)-oxide nanoparticles of the birnessite type comprising 50-100 Mn ions per PSII. Having shown that even today’s photosystem-II can form birnessite-type oxide particles efficiently, we propose an evolutionary scenario, which involves Mn-oxide production by ancestral photosystems, later followed by down-sizing of protein-bound Mn-oxide nanoparticles to finally yield today’s Mn_4_CaO_5_ cluster of photosynthetic water oxidation.

## Introduction

Nature’s invention of photosynthetic water oxidation about three billion years ago (or even earlier^1^) was a breakpoint in Earth’s history because it changed the previously anoxic atmosphere to today’s composition with ~21 % O_2_, practically depleting the oceans of ferrous iron and divalent manganese due to metal-oxide precipitation^2,3^. Water oxidation is catalyzed by a unique bioinorganic cofactor, denoted Mn_4_CaO_5_ according to its oxo-bridged metal core, which is bound to amino acids of the proteins of photosystem II (PSII) in the thylakoid membrane (Fig. 1)^4–8^. This catalyst originally developed in (prokaryotic) cyanobacteria, which were later incorporated by endosymbiosis into the ancestor of the (eukaryotic) cells of algae and plants to yield the chloroplast organelles^9^. The central PSII proteins as well as Mn_4_CaO_5_ (and its main catalytic performance features) are strictly conserved among photosynthetic organisms^10^.

**Figure 1:**
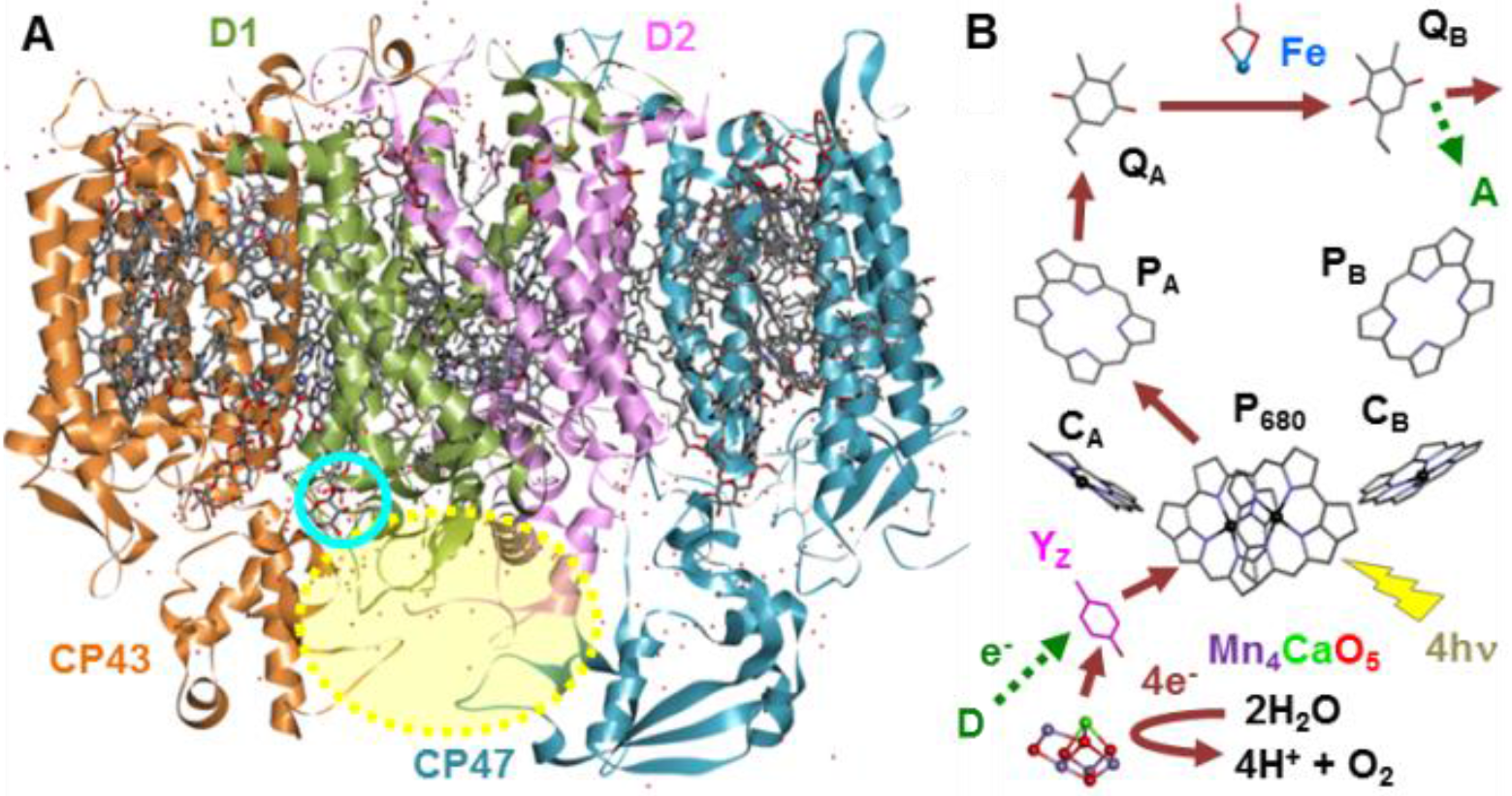
Structure and function of photosystem II. (A) Crystal structure of native plant PSII (PDB entry 5XNL^11^). The central membrane-integral subunits with cofactors of a PSII monomer are shown (cyan circle = Mn_4_CaO_5_ position). Omitting extrinsic subunits (18, 24, 33 kDa; absent in our Mn-depleted PSII) exposes a cavity (yellow). (B) Cofactors (truncated structures) for light-driven electron transfer in D1/D2 (Y_Z_, tyrosine electron acceptor from Mn_4_CaO_5_; P_680_, primary donor chlorophyll (chl) dimer; C_A,B_, accessory chl; P_A.B_, pheophytins; Q_A,B_, acceptor quinones; Fe, non-heme iron with bicarbonate ligand). Green/dark-red arrows: electron transfer paths upon photo-excitation (*hv*) of O_2_-evolving or Mn-depleted PSII (D/A, external electron donor/acceptor).

Native PSII operates as a light-driven oxidoreductase (Fig. 1). Upon sequential excitation with four visible-light photons, four electrons from two bound water molecules are transferred from Mn_4_CaO_5_ to a redox-active tyrosine (Y_Z_) at the donor side and then via a cofactor chain to terminal quinone acceptors at the stromal side so that two reduced quinols as well as O_2_ and four protons are released during each catalytic water oxidation cycle^5,7,12–14^. Starting from a Mn(III)_3_Mn(IV)Y_Z_ state, the catalytic cycle involves alternate electron and proton abstraction to reach a Mn(IV)_4_Y_Z^ox^_ state followed by (concomitant) Mn re-reduction, O-O bond formation and O_2_ release (Fig. 1)^15^. The exceptionally efficient Mn_4_CaO_5_ catalyst has inspired development of synthetic water-oxidizing materials^16–18^. Among the wealth of findings on water oxidation by Mn-based catalysts, here the following two results are of particular importance: (i) Self-assembly of the Mn(III/IV)_4_CaO_5_ core in PSII is a light-driven process, involving step-wise oxidation of four solvent Mn^2+^ ions by Y_z^ox^_ coupled to electron transfer to the quinones^19,20^. (ii) Many amorphous Mn oxides of the birnessite type show significant water oxidation activity and share structural as well as functional features with the Mn_4_CaO_5_ core of the biological catalyst^21–25^.

The evolutionary route towards the present water oxidation catalyst in PSII is much debated^2,26–31^. It has been hypothesized that before the evolution of oxygenic photosynthesis an ancestral photosystem eveloped the capability for light-driven oxidation of dissolved Mn^2+^ ions towards the Mn(III/IV) level, thereby providing the reducing equivalents (electrons) needed for primary biomass formation by CO_2_ fixation^32,33^. Noteworthy, the process of continuous Mn^2+^ oxidation is chemically not trivial, because suitable redox potentials alone are insuficient. Because solitary Mn(III/IV) ions are not stable in aqueous solution, the ability of the photosystem to stabilize high-valent Mn ions by efficient formation of extended Mn (oxide) structures is pivotal. In the present study, clear experimental evidence is provided that today’s PSII, depleted of its native Mn_4_CaO_5_ complex and the membrane-extrinsic polypeptides, can form a Mn(III/IV) oxide of the birnessite type. Aside from the implications for biological evolution, photosynthetic Mn oxide formation has significance in the context of recent hypotheses to account for geologic Mn deposits, for example from the early Paleoproterozoic in South Africa^2,33^.

## Results

### Spinach photosystems depleted of Mn_4_CaO_5_ and extrinsic polypeptides

Figure 1 shows the arrangement of protein subunits and cofactors in PSII. A recent crystallographic study has revealed that the metal-binding amino acids are similarly arranged in PSII with or without Mn_4_CaO_5_, with the voids in the Mn-depleted photosystem being filled by water molecules^34^. Only in the absence of the Mn-stabilizing extrinsic proteins, sufficient room for incorporation of a Mn oxide nanoparticle into the PSII structure may exist (Fig. 1). Therefore, we explored the ability of purified PSII, depleted of Mn_4_CaO_5_ and the extrinsic proteins, to form Mn-oxide species *in vitro*. PSII-enriched membrane particles were prepared from spinach^35^ and Mn depletion was achieved using an established protocol (see Supporting Information)^36^. The resulting PSII preparation was inactive in light-driven O_2_-evolution and Mn was practically undetectable, i.e., less than 0.2 Mn ions per PSII were found (Table 1). Concomitantly with Mn depletion, the three proteins bound to the lumenal side of plant PSII (the so-called extrinsic proteins of about 18 kDa, 23 kDa, and 33 kDa)^37,38^ were removed, as revealed by polarography, TXRF metal quantification, and gel electrophoresis (Figs. S1, S2; Tables 1, S1).

**Table 1:**
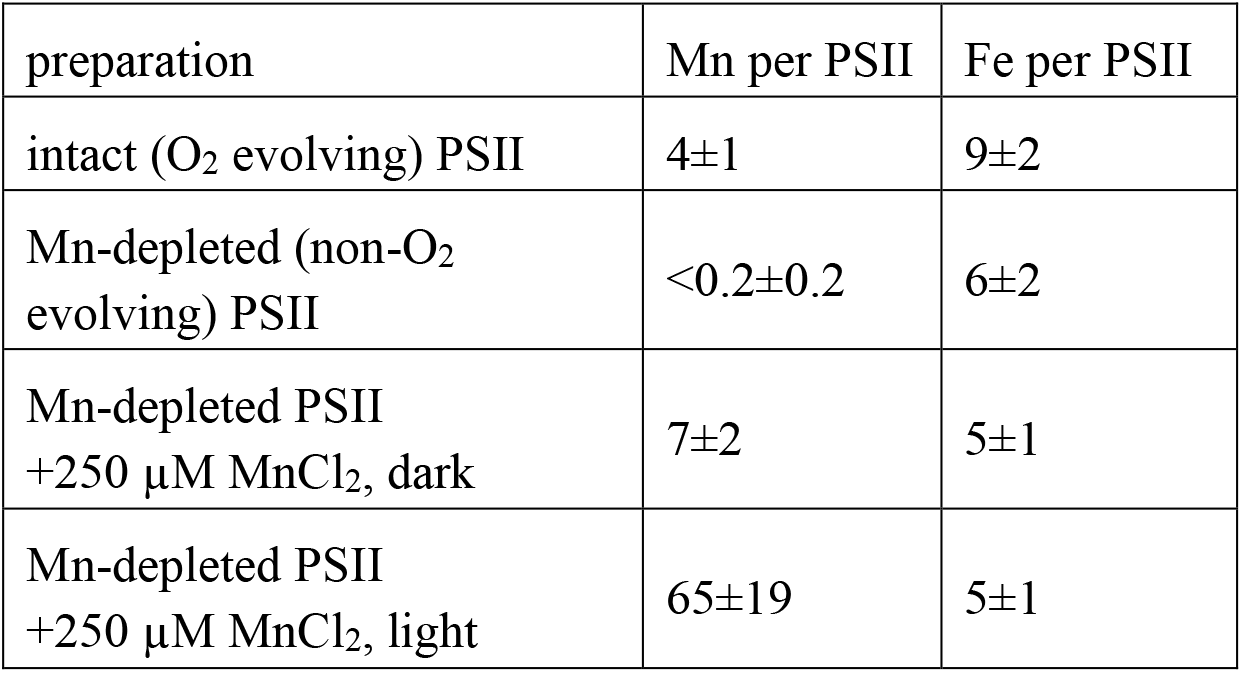
Metal content of PSII preparations (for details, see Table S1).

### UV-vis spectra monitor PSII electron transfer

Optical absorption spectroscopy was employed for time-resolved tracking of PSII redox chemistry (Figs. 2-4 and S3-S9). DCPIP (2,6-dichlorphenol-indophenol) was added as an artificial electron acceptor for PSII that allows facile optical monitoring of light-driven electron flow^39^. Oxidized DCPIP (DCPIP^ox^) at pH ≥ 7 shows strong absorption at 604 nm and its bright blue color vanishes upon reduction (Figs. 2, 3, see also Fig. S5)^40^. We explored the ability of DCPIP^ox^ to support and simultaneously probe PSII electron transfer by recording absorption spectra for increasing illumination periods. For intact PSII with a native Mn_4_CaO_5_ cluster, rapid and essentially complete DCPIP reduction was observed (Figs. S5, S6). Mn-depleted PSII showed a completely different electron transfer behavior (Figs. 3, 4 and S7, S9). By adding a Mn salt (MnCl_2_), we investigated hexaquo-Mn^2+^ ions as an exogenous electron donor to PSII. Mn^2+^ in solution is completely colorless, i.e., it does not absorb in the 300-900 nm region. In the dark in the presence of DCPIP^ox^ and Mn^2+^ ions or upon illumination in the absence of Mn^2+^, we did not observe any major spectral change of the PSII suspension (aside from minor bleaching of PSII chlorophyll due to oxidative damage^41^). No absorption changes accountable to DCPIP (i.e., due to its reduction) under illumination were observed in the presence of MnCl_2_ with simultaneous absence of PSII (Fig. S8). These experiments verify for the Mn-depleted PSII: electron transfer towards DCPIP requires both, visible light to drive the PSII electron transfer reactions and Mn^2+^ ions that can serve as a donor in the light-induced electron transfer.

**Figure 2:**
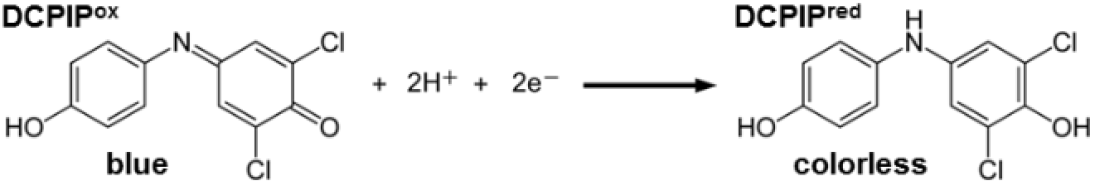
Oxidized and reduced DCPIP (2,6-dichlorphenol-indophenol).

**Figure 3:**
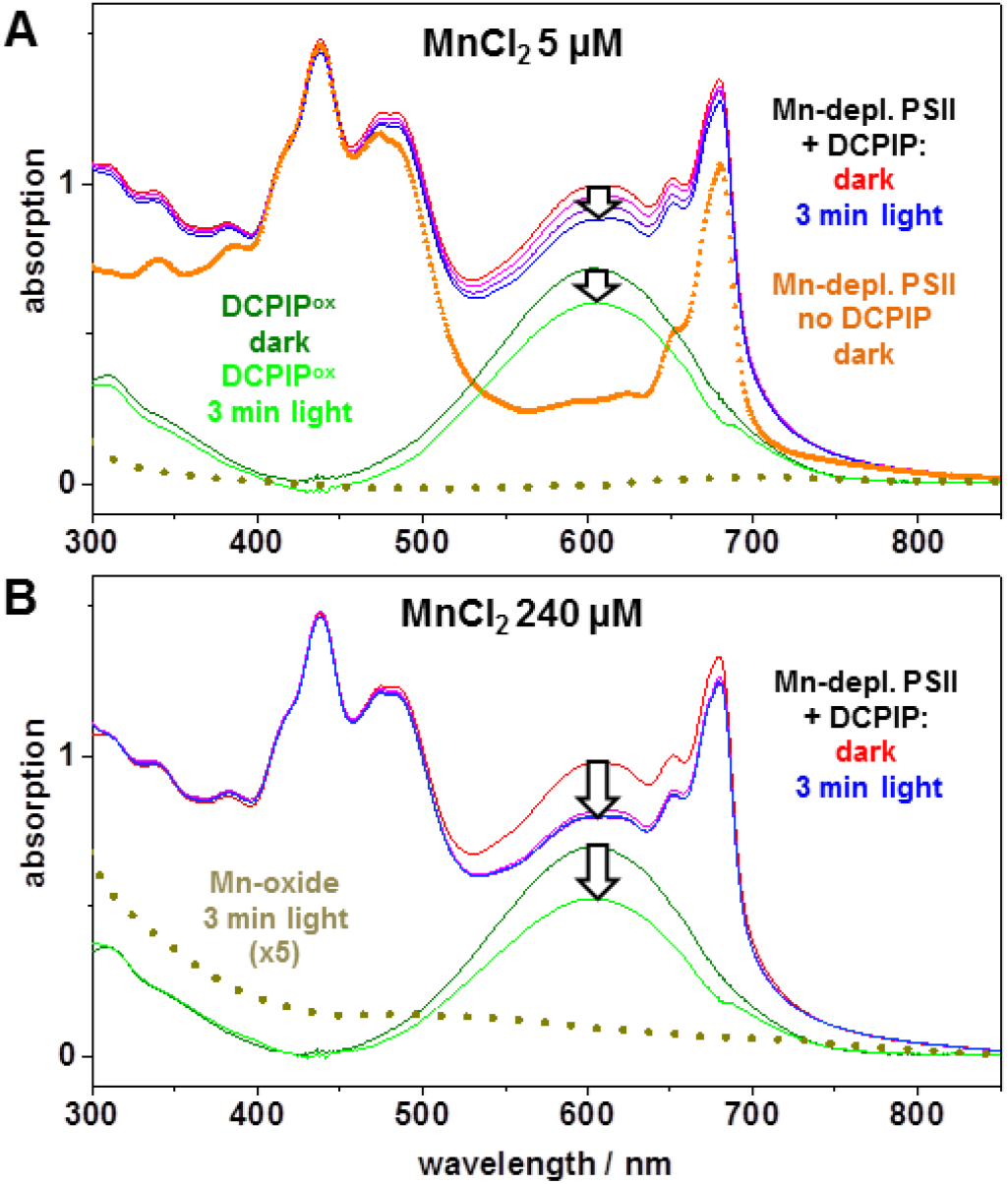
Light-driven redox reactions in Mn-depleted PSII tracked by optical spectroscopy. (A) and (B) show UV-vis absorption spectra. The orange line (triangles) represents a suspension of Mn-depleted PSII before addition of DCPIP (60 μM) serving as artificial electron acceptor (buffer conditions: 1 M glycine-betaine, 15 mM NaCl, 5 mM CaCl_2_, 5 mM MgCl_2_, 25 mM MES buffer, pH 7; for further details see SI). Then DCPIP^ox^ and either 5 μM MnCl_2_ (in A) or 240 μM MnCl_2_ (in B) were added, followed by continuous white-light illumination (1000 μE m^-2^ s^-1^) of the PSII suspensions and collection of spectra (one spectrum per minute; immediately prior to illumination, red lines, and after 3 min light, blue lines). The green and light-green lines correspond to the spectral contributions of DCPIP^ox^ to the dark and 3-min light spectra; the amplitude decrease at 604 nm represents the loss of DCPIP^ox^ due to its reduction by PSII (arrows). The dark-yellow dotted spectra are assignable to a Mn-oxide, as verified by X-ray absorption spectroscopy (spectra obtained by weighted spectral deconvolution, see caption of Fig. S10, and scaled by a factor of 5, for clarity). Note that significant spectral changes due to Mn-oxide formation were only observed with 240 μM MnCl_2_ (in B), but not with 5 μM MnCl_2_ (in A).

### Electron donation by Mn^2+^ ions

At low MnCl_2_ concentration (5 μM), a linear decrease in the amount of DCPIP^ox^ indicates a small and constant rate of electron donation at the PSII donor side (Figs. 3, 4), which is only about 8% of the level reached in intact PSII. It may be assignable to PSII centers that either acquire an alternative minor reactivity or support light-driven oxygen evolution at exceedingly low rate. This slow electron transfer is not visible at higher MnCl_2_ concentrations suggesting that at high MnCl_2_ concentrations, the PSII centers do not acquire a metal complex that supports continuous low-rate electron transfer. Therefore we consider this phenomenon, albeit of clear interest for future investigation, irrelevant for the analysis of oxide formation herein observed at higher MnCl_2_ concentrations. For increasing MnCl_2_ concentrations, a clearly more rapid phase of DCPIP^ox^ reduction grew in (Fig. 3, arrows; Figs. 4 and S7). Its amplitude saturated at 240 μM MnCl_2_ and indicates reduction of ~17 % (~10 μM) of the initial DCPIP^ox^ (60 μM), which corresponds to up to 200 transferred electrons per PSII. Based on the Mn concentration dependence and the results presented in the following, we can assign this rapid DCPIP reduction phase to oxidation of Mn^2+^ ions and formation of high-valent Mn(III/IV) oxide particles. At 240 μM MnCl_2_, DCPIP reduction is completed after about 1 min of illumination likely indicating that further electron donation to the PSII donor side (and thus further DCPIP reduction) was impaired after the formation of a Mn oxide particle at the PSII donor side that blocks access of further Mn^2+^ ions to the oxidant, which is the tyrosine (YZ) radical (see Fig. 1).

**Figure 4:**
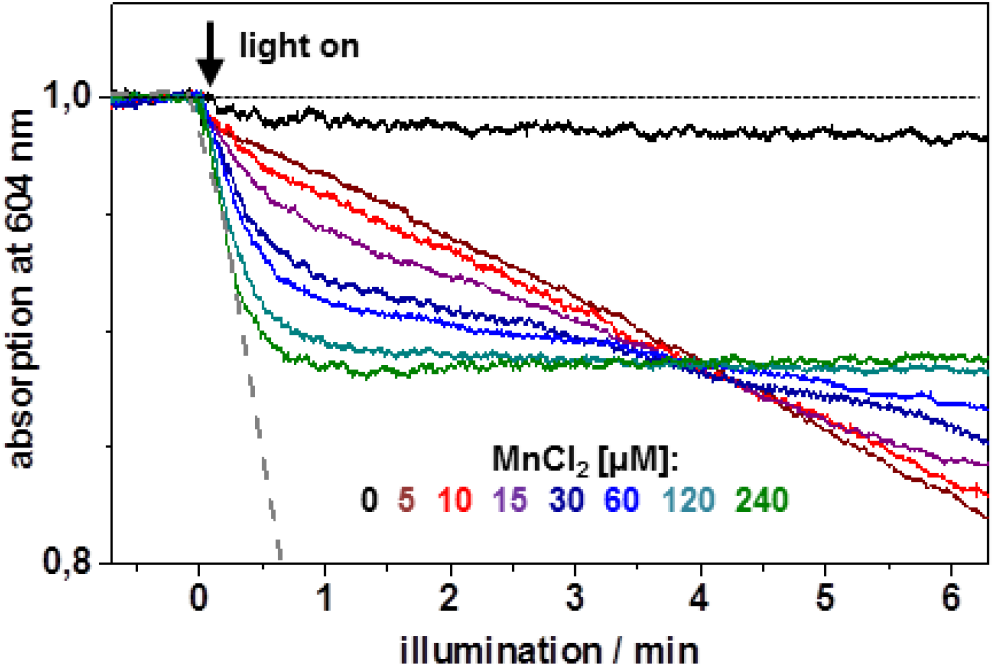
Kinetics of light-dependent electron flow in Mn-depleted PSII at various concentrations of solvated Mn^2+^ ions. The decrease in absorption at 604 nm monitors the light-driven reduction of the artificial electron acceptor (60 μM DCPIP^ox^, pH 7) due to the oxidation of Mn^2+^ ions by PSII. The grey dashed line illustrates the rate of DCPIP reduction by fully intact PSII under similar conditions (Fig. S6).

### UV-vis spectra point towards Mn-oxide formation

To search for evidence of Mn oxide formation, informative absorption difference spectra of Mn-depleted PSII before and after illumination were calculated (Figs. 3 and S7). For 60 μM MnCl_2_, after completion of rapid DCPIP^ox^ reduction (3 min), there was a broad absorption increase (ranging from 350-700 nm), which is similar to the wide-range absorption of Mn oxides^25^. For higher concentrations of MnCl_2_, the absorption assigned to Mn oxides gained strength and became maximal at 240 μM MnCl_2_ (Fig. 3). Using alternative electron acceptors (Fig. S9), similar or even higher Mn-oxide amounts were detected with DCBQ (2,5-dichloro-1,4-benzoquinone) or PPBQ (phenyl-p-benzoquinone), resembling the native quinone acceptor (QB), but the slow (hydrophilic) acceptor ferricyanide (K_3_Fe^III^(CN)_6_) did not yield significant Mn-oxide formation.

### PSII with 50-100 bound Mn ions prepared for analysis by X-ray spectroscopy

To investigate the Mn^2+^ oxidation products and identify their atomic structure, we employed X-ray absorption spectroscopy (XAS) at the Mn K-edge (Figs. 5 and S10, S11). Mn-depleted PSII was illuminated for 3 min with 240 μM MnCl_2_ and 60 μM PPBQ^ox^ (pH 7.5), the reaction was terminated by rapid sample cooling in the dark, and the PSII membranes were pelleted by centrifugation and then transferred to XAS sample holders, followed by freezing in liquid nitrogen and later collection of X-ray spectra at 20 K (see SI). The metal content was determined by X-ray fluorescence analytics (Tables 1, S1), revealing 65±19 Mn ions per initially Mn-depleted PSII after illumination in the presence of 240 μM MnCl_2_. The calcium content in the PSII-formed Mn oxide could not be reliably determined because CaCl_2_ was present in the buffer and Ca is known to bind unspecifically to the used PSII membrane particle preparation (Table S1)^42^.

**Figure 5:**
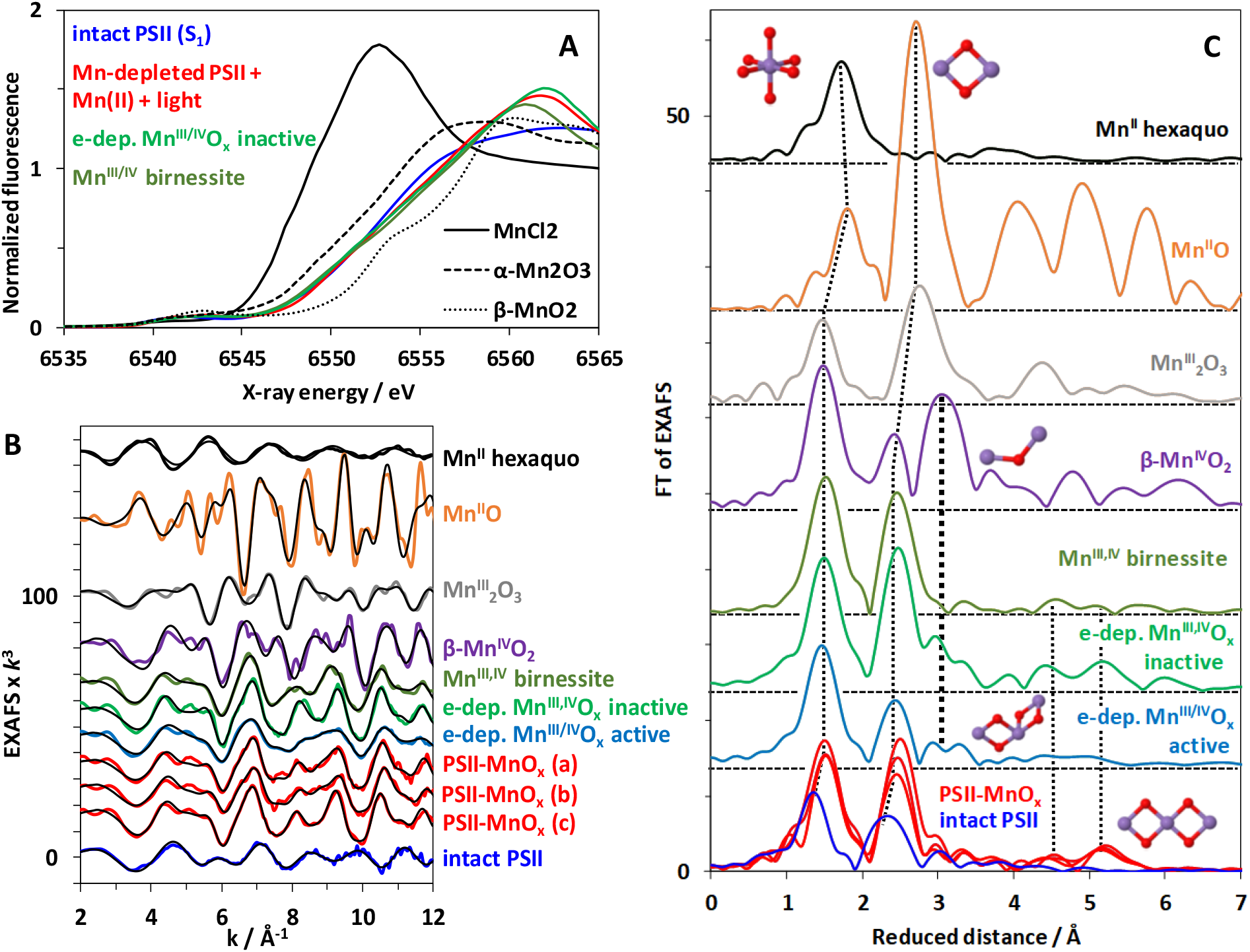
X-ray absorption spectroscopy evidencing PSII-bound Mn oxide of the birnessite-type. (**A**) XANES spectra of the native PSII with its active-site Mn(III)2Mn(IV)2CaO_5_ cluster (blue) and Mn-depleted PSII after light-driven formation of Mn oxide (red) are compared to the spectra of Mn^2+^ in solution, electrodeposited Mn-oxide (green), synthetic birnessite (dark green) and further Mn oxides (black lines). (**B**) The corresponding EXAFS spectra in *k*-space as in A as well as spectra of further Mn compounds. (**C**) Fourier-transformed (FT) EXAFS spectra (as in B) of: Mn^2+^ ions in solution, indicated reference Mn oxides, the Mn_4_CaO_5_ cluster in intact PSII, and Mn-depleted PSII after illumination in the presence of Mn^2+^. Structural motifs corresponding to the individual peaks are schematically shown; FT peaks relating to the same structural motif are connected by dotted lines. Mn-depleted PSII particles (20 μg mL^-1^ chlorophyll) were illuminated for 3 min (4000 μE m^-2^ s^-1^) in the presence of 240 μM MnCl_2_ and 60 μM PPBQ (Fig. S9; buffer conditions: 1 M glycine-betaine, 15 mM NaCl, 5 mM CaCl_2_, 5 mM MgCl_2_, 25 mM MES buffer, pH 8) and subsequently collected by centrifugation. (The spectra shown in A as a red line represents the mean of the three spectra from individual samples shown in B and C as red lines as obtained after subtraction of a 5 % aqueous Mn^2+^ contribution. See Table S2 for details and further data.)

### X-ray spectroscopy reveals extended Mn(III/IV) oxides

The shape of the XANES (X-ray absorption near-edge structure) pronouncedly differed from hexaquo-Mn^2+^, micro-crystalline Mn oxides (Mn^III^_2_O_3_, Mn^II,III^_3_O_4_, β-Mn^IV^O_2_), and native PSII, but was similar to layered Mn(III,IV) oxides denoted as birnessite^43–45^ (Fig. 5A). The K-edge energy indicated a mean redox level of about +3.5, suggesting equal amounts of Mn(III) and Mn(IV) ions (Figs. 5A, S10). EXAFS (extended X-ray absorption fine structure) analysis revealed the atomic structure of the PSII-bound Mn-oxide (Figs. 5B, S11; Table S2). The EXAFS of Mn-depleted PSII with bound Mn oxide closely resembled birnessite (44) in showing a similar main Mn-O bond length (~1.90 Å), minor longer Mn-O bond length contributions (~2.30 Å, assignable to Jahn-Teller elongated Mn-O distances of Mn^III^ ions) as well as similar main and minor Mn-Mn distances (~2.88 Å, ~3.45 Å). Also, longer Mn-Mn distances (~5.00 Å, ~5.54 Å) were similar. On the other hand, the EXAFS spectra differ clearly from Mn(III)_2_O_3_ and β-Mn(IV)O_2_. The metrical parameters from EXAFS simulations are in good agreement with earlier data for the here studied and related Mn oxide species of the birnessite-type^21,22,25,46,47^. We note that the long-range order in the oxide particles produced by PSII even exceeds that of the herein used reference oxides of the birnessite-type, as indicated by the magnitudes of the Fourier peaks assignable to the 2.87 and 5.54 Å distances, verifying formation of a comparably extended and well-ordered Mn oxide. Notably, according to the similar XAS spectra, a similar birnessite-type Mn(III,IV)-oxide was formed both in the presence and absence of CaCl_2_ in the illumination buffer (Fig. S12).

## Discussion

### Mn oxide formation by PSII

We have obtained the first direct experimental evidence that PSII devoid of Mn_4_CaO_5_ is capable of forming Mn(III,IV)-oxide particles of the birnessite type by light-driven oxidation of Mn^2+^ ions. Light-driven Mn^2+^ oxidation also can promote self-assembly of functional Mn_4_CaO_5_, which is a comparably inefficient (low quantum yield) low-light process denoted as photoactivation^19,20^. Cheniae et al. investigated photoactivation and observed light-driven binding of about 18 membrane-bound Mn ions per PSII if Ca ions were excluded from the photoassembly buffer^48^, whereas we here observe binding of clearly more Mn ions (65±19 Mn ions, Table 1), irrespective of the absence or presence of Ca ions at moderate concentration (5 mM) in the photoassembly buffer (Figure S12). The presence of 5 mM CaCl_2_ allows for photoactivation, although higher concentration are required for optimal photoactivation yield^48,49^. The about 20 times higher light intensities we used likely promoted the oxidation and binding of numerous Mn ions at the expense of formation of a single native Mn_4_CaO_5_ cluster, because the latter requires low light intensities presumably due to the presence of a slow ‘dark rearrangement’ step for assembly^19^

Since Cheniae’s work, it had remained an open question in what form a larger number of Mn ions can bind to PSII membrane particles. Coordination of individual high-valent Mn ions to protein groups is one possibility (as often observed for divalent cations and trivalent Fe ions); the formation of extended protein-bound Mn-oxides nanoparticles is another possibility. Under our high-light conditions, Mn(III,IV)-oxide formation clearly is dominant. The Mn-oxide cluster size seems to be limited to around 100 Mn ions, which may correspond to a nanoparticle of about 20 Å in diameter (Fig. 6A). Such a particle may well be adopted in the PSII cavity that becomes solvent-exposed upon removal of the extrinsic protein subunits (Fig. 1). These subunits are evolutionarily younger than the PSII core proteins^50^ and are absent in related anoxygenic photosystems^51^. Thus, it is well conceivable that an early PSII ancestor would have lacked these extrinsic proteins and therefore could accommodate a Mn-oxide nanoparticle. Furthermore, an ancient autotroph, capable of exploiting Mn^2+^ as a metabolic reductant^32,33^, would be expected to be configured so that the donor side of the early PSII ancestor would be exposed to the environment as opposed to being sequestered within the lumen of modern thylakoids. In this context, the cyanobacterium *Gloeobacter violaceous* provides an interesting example.*Gloeobacter* occupies a basal phylogenetic position and evolved before the appearance of thylakoids. It possesses photosynthetic reaction centers that are located in the cytoplasmic membrane with the oxidizing domain of photosystem II facing the periplasmic space and thus the exterior of the cell^52^. Thus *Gloeobacter* provides an example of how a primordial reaction center might have been arranged to facilitate the photochemical utilization of Mn^2+^ as a reductant source, as originally proposed by Zubay^32^.

**Figure 6:**
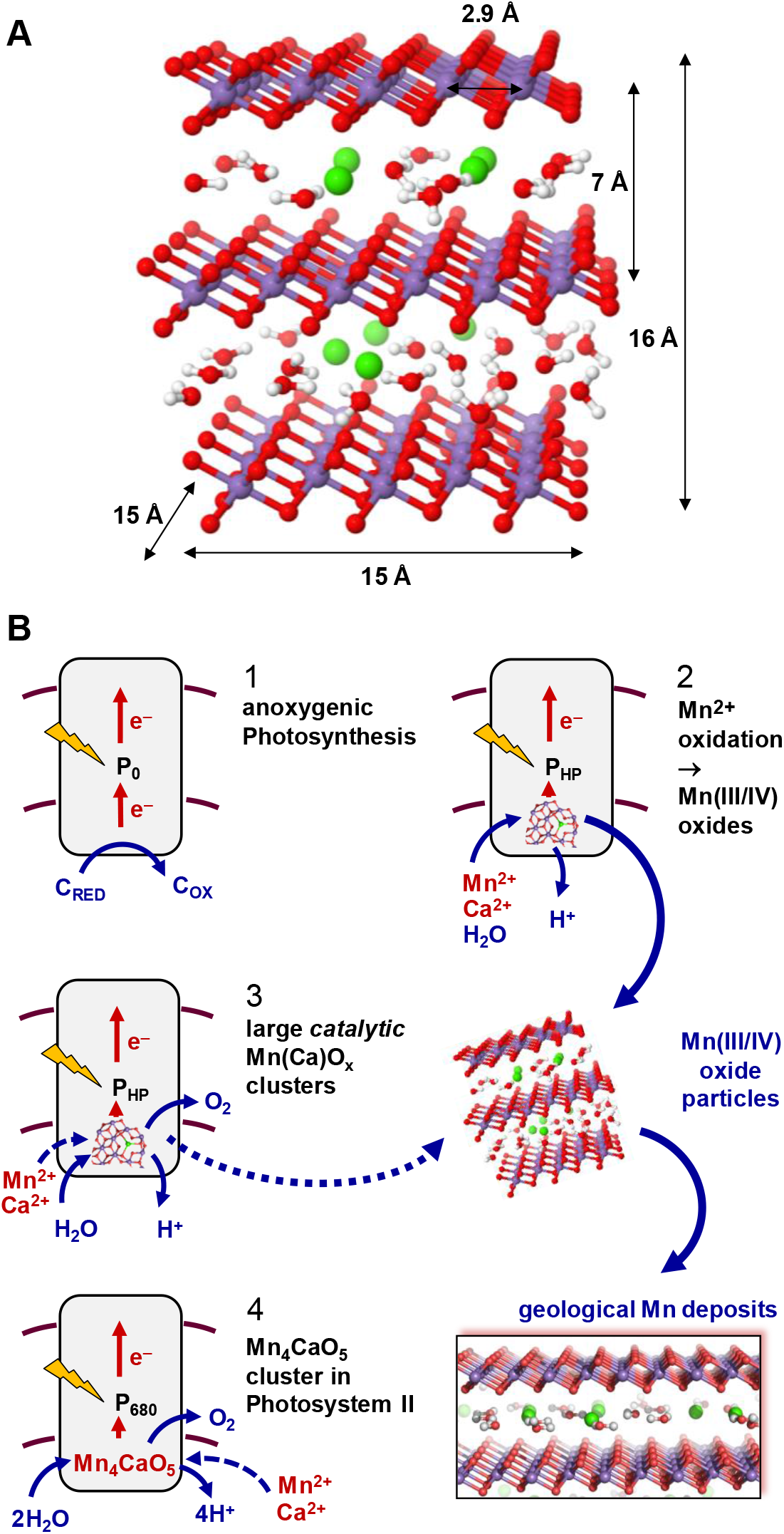
Evolution of oxygenic photosynthesis and relation to geological Mn-oxide deposits. (**A**) Structural model of a birnessite fragment with 64 Mn ions (based on the atomic coordinates in ref.^61^, protons were added for illustration only). Atom color coding: violet, Mn(III/IV) ions; red, O or OH; green, Ca^2+^; grey, H. (**B**) Proposed sequence of evolutionary events. (1) Starting from an anoxygenic photosystem with a low-potential primary donor (P_0_) in a marine cyanobacterial ancestor, (2) a higher donor potential (P_HP_) facilitated Mn^2+^ oxidation^62^ to generate protein-bound Mn(III,IV)-oxide clusters, which upon sedimentation contributed to geological Mn-oxide deposits^33^. (3) The PSII-bound clusters developed into primordial water-oxidizing and O_2_-evolving catalytic complexes, which, (4) due to a dedicated binding site, were down-sized to the present Mn_4_CaO_5_ catalyst.

### Relation to water-oxidizing synthetic Mn oxides

Birnessite and buserite are layered, typically non-crystalline metal-oxides with sheets of edge-sharing MnO_6_ octahedra (which corresponds to di-μ-oxo bridging between neighboring Mn ions) separated by water and cations, e.g., Na^+^ or Ca^2+^, in the interlayer space^43,44^ Birnessite and buserite differ regarding the number of water-cation layers in between two oxide layers (one in birnessite, two in buserite), but share the same fundamental structure of the Mn(III,IV) oxide layers and thus are often jointly denoted as birnessite-type Mn oxides. Noteworthy, a Mn oxide denoted as ranciéite is isostructural to birnessite and contains Mn and Ca ions at approximately the same 4:1 stoichiometry as present in the Mn_4_CaO_5_ cluster of PSII^53^, suggesting a possible relation^27,54–56^. Birnessite-type Mn oxides are a major component of Mn-oxide ocean nodules43and biogenic Mn oxides^57^. Their diagenetic reductive conversion to Mn-bearing carbonates, on geological time scales, may explain the early Mn deposits reported by Johnson et al.^33^. Notably, by electrodeposition and other synthesis protocols, Mn(III/IV) oxides of the birnessite type can be formed that are either active or largely inactive in water oxidation, depending on their atomic structure^21,22,25,46,58,59^. These Mn(III/IV) oxides share key features with the Mn_4_CaO_5_ cluster of the biological catalyst, including joint structural motifs and facile oxidation state changes during catalytic operation^24^. The presence of Ca ions is especially favorable for water oxidation activity by synthetic manganese oxides, pointing towards similar water-oxidation mechanisms in the synthetic oxides and the biological Mn_4_CaO_5_ cluster of PSII^21,22,25,60^. Regarding their high degree of structural order, the Mn oxide particles formed by PSII resemble electrodeposited Mn oxides that are able to undergo Mn(III)—Mn(IV) redox transitions, but exhibit low electrochemical water oxidation activity^24^. The structural characteristics that have been identified for transforming a largely inactive Mn oxide into an oxide with sizeable water-oxidation activity^24^ are apparently lacking in the Mn oxide particles formed by PSII, which may explain the absence of detectable water-oxidation activity by the herein investigated PSII-bound Mn oxide particle.

### Mechanism of Mn-oxide formation

The basic biochemical mechanism of the here described light-induced Mn-oxide formation likely involves initial binding of Mn^2+^ ions followed by Mn oxidation and stabilization of the oxidized Mn(III/IV) ions by di-μ-oxo bridging, in analogy to both the photoassembly process of today’s Mn_4_CaO_5_ cluster^19,20^ and the “oxidative self-assembly” process in electrodeposition of non-biological birnessite-type Mn oxides^25,46^. The formation of extended oxide particles likely involves a nucleation-and-growth mechanism. In the photosystem, the initial site of Mn^2+^ binding and formation of an oxide nucleus likely is provided by carboxylate and possibly imidazole sidechains of protein residues followed by an oxide growth that does not require further ligating residues.

### Oxide-incorporation hypothesis on the evolution of today’s Mn_4_CaO_5_ cluster

Various hypotheses on the origin of the Mn_4_CaO_5_ cluster have been proposed. According to Raymond and Blankenship the interaction of an anoxygenic PSII with a manganese catalase, utilizing hydrogen peroxide as the source of electrons, led to the formation of today’s Mn_4_CaO_5_ cluster^28^, without invoking Mn oxides. Dismukes and coworkers developed hypotheses on the evolution of oxygenic photosynthesis by focusing on the inorganic chemistry of Mn and bicarbonate^63^; their analyses could complement the herein developed ideas on the evolutionary role of Mn oxides in the future. In 2001, Russell and Hall developed their influential Mn-oxide incorporation hypothesis^54^. They suggested that a “ready-made”cluster must have been co-opted whole by a (mutant?) protein..”^55,56^. Russell and Hall specifically proposed that dissolved Mn^2+^ ions were photo-oxidized at extremely short wavelengths64to colloidal clusters of [CaMn_4_O_9_ ·3H_2_O], which are closely related to the birnessite-type Mn oxide denoted as rancicéite. Incorporation of this or a similar “ready-made” Mn_4_Ca species into a PSII ancestor would have led to the Mn_4_CaO_5_ cluster of today’s PSII. This hypothesis is in line with analyses of Yachandra and Sauer who systematically compared the structural relations between various Mn oxides and the biological metal complex, revealing intriguing similarities^27^ (for further discussion, see SI Appendix – background text).

### Alternative hypothesis on evolution of the Mn_4_CaO_5_ cluster

We see a close relation between inorganic Mn oxides and today’s Mn_4_CaO_5_ cluster of PSII that differs distinctively from the Mn-oxide incorporation hypotheses outlined above. In our study, facile formation of birnessite-type Mn oxide particles by PSII is reported. They (i) share structural motifs with the biological cluster in PSII^10,65^ and biogenic Mn oxides in general^57,66^ and (ii) resemble synthetic Mn oxides closely that have been investigated as synthetic catalyst materials^24^. On these grounds we propose a scenario illustrated in Figure 6: Rather than Mn oxide incorporation, Mn oxide nanoparticles were formed by an evolutionary precursor of PSII, *inter alia* enabling the formation of early geological Mn oxide deposits. Initially, dissolved Mn^2+^ ions served as a source of reducing equivalents eventually needed for CO_2_ reduction, as has been suggested first by Zubay^32^ and later by others^31,33^. At a later stage, down-sized oxide particles developed into today’s water-oxidizing Mn_4_CaO_5_ cluster. Are there evolutionary relicts that may support the above hypothesis? Extensive studies on the diversity of the PSII reaction center protein D1 have revealed several atypical variants that can be distinguished phylogenetically^67,68^. These early evolved forms lack many residues needed for the binding of today’s Mn_4_CaO_5_ cluster and could relate to ancient Mn oxide forming photosystems, even though today they might play other physiological roles (e.g., in the synthesis of chlorophyll *f*^69^).

### Summary of potential evolutionary implications

The ability for light-driven Mn oxide formation by an ancient photosystem represents an important touchstone for evaluation of three interrelated hypotheses that each addresses a remarkable facet of the evolution of the Earth’s biosphere and geosphere:

i. The ability for direct and facile photosynthetic formation of stable Mn(III/IV)-oxide particles supports that early Mn deposits^33^ resulted directly from photosynthetic activity.
ii. Structural and functional similarities between water-oxidizing synthetic Mn oxides and the here described Mn-oxide formation by PSII suggests that in the evolution of PSII, there may have been a transition from extended Mn-oxide nanoparticles towards the Mn_4_CaO_5_ cluster of today’s PSII, as illustrated by Figure 6.
iii. An early quasi-respiratory cycle has been proposed that involves formation of Mn(III/IV) oxide particles followed by utilization of the oxidizing equivalents stored in the Mn oxide for an efficient quasi-respiratory activity in the Archean or early Paleoproterozoic, when the Earth’s atmosphere had been essentially O_2_-free, as detailed in^66^.

By showing that today’s PSII can form birnessite-type Mn-oxide particles efficiently, even without any specific protein subunits that would support Mn-oxide formation, the general biochemical feasibility is verified. This finding renders it highly likely that similarly also an ancient photosystem, the PSII ancestor, had the ability for light-driven formation of Mn-oxides from hexaquo Mn^2+^ ions. In conclusion, we believe that our successful demonstration of photosynthetic formation of Mn(III/IV)-oxide particles provides relevant support for the above three hypotheses.

## Methods

### Preparation of PSII membrane particles

Native PSII-enriched thylakoid membrane particles were prepared from fresh market spinach following our established procedures^35^. Their typical O_2_-evolution activity (as determined by polarography with a Clark-type electrode at 27 °C) was ~1200 μmol O_2_ mg^-1^ chlorophyll h^-1^, which proved the full integrity of the PSII proteins and the water-oxidizing Mn_4_CaO_5_ complex. We have shown earlier that this type of PSII preparation contains ~200 chlorophyll molecules per PSII reaction center^70,71^. When kept for prolonged time periods in the dark, the Mn_4_CaO_5_ complex is synchronized in the S_1_ state of its catalytic cycle, which is established to represent a Mn(III)_2_Mn(IV)_2_ oxidation state^72,73^.

### Mn-depletion of PSII

Removal of Mn_4_CaO_5_ and of the three extrinsic proteins of PSII (18, 24, 33 kDa) was carried out using a literature procedure and evaluated using metal quantification and gel electrophoresis (see below)^36^. PSII membranes were dissolved at 200 μg chlorophyll mL^-1^ in a high-salt buffer (30 mL) containing 20 mM TEMED (N,N,N’,N’-tetramethylethylenediamine) as a reductant for the PSII-bound Mn(III,IV) ions, 20 mM MES (2-(N-morpholino)ethane-sulfonic-acid) buffer (pH 6.5), and a high salt concentration (500 mM MgCl_2_) and incubated in the dark on ice for 10 min. PSII membranes were pelleted by centrifugation (Sorvall RC26, 12 min, 50000 g, 4 °C), the pellet was 3-times washed by dissolution in a buffer (30 mL) containing 35 mM NaCl and 20 mM TRIS (tris(hydroxymethyl)aminomethane) buffer (pH 9.0) and pelleting by centrifugation as above, and the final pellet of Mn-depleted PSII membranes was dissolved at ~1 mg chlorophyll mL^-1^in a buffer containing 1 M glycine-betaine, 15 mM NaCl, 5 mM CaCl_2_, 5 mM MgCl_2_, and 25 mM MES buffer (pH 6.3). The PSII preparations (~2 mg chlorophyll mL^-1^) were thoroughly homogenized by gentle brushing and frozen in liquid nitrogen for the spectroscopic experiments. The Mn-depleted PSII showed zero O_2_-evolution activity as revealed by polarography.

### Total reflection X-ray fluorescence (TXRF) analysis

X-ray emission spectra were recorded on a Picofox instrument (Bruker) and metal contents of PSII samples were determined from the data using the (fit) routines available with the spectrometer74PSII membranes were adjusted to a chlorophyll concentration of 1-2 mM and to a 20 μL aliquot, a gallium concentration standard (1 mg mL^-1^, 20 μL; Sigma-Aldrich) was added, and samples were homogenized by brief sonication (see Fig. S1). A 5 μL aliquot of the samples was pipetted on clean quartz discs for TXRF, dried on a heating plate, loaded into the spectrometer, and TXRF spectra were recorded within 10-30 min. At least 3 repetitions of each sample and 3 independently prepared samples of each PSII preparation were analyzed.

### Optical absorption spectroscopy and illumination procedures

For the optical absorption spectroscopy experiments, stock suspensions of the PSII preparations were diluted at 20 μg chlorophyll mL^-1^ (~0.1 μM PSII centers) in a buffer (3 mL) containing 1 M glycine-betaine, 15 mM NaCl, 5 mM CaCl_2_, 5 mM MgCl_2_, and 25 mM MES buffer (pH 6.3-8.5) and reactants (oxidized 2,6-dichlorophenol-indophenol = DCPIP^ox^ from Fluka, other electron acceptors as in Fig. S9, MnCl_2_) were added at indicated concentrations. Optical absorption spectra of the samples in a 300-900 nm range were recorded within about 10 s at given time intervals (about 0.3-1.0 min) in a 3 mL quartz cuvette (Helma QS1000, 1 cm pathlength) using a Cary 60 spectrometer (Agilent). Alternatively, time traces of absorption were recorded at selected wavelengths (i.e. 604 nm to monitor DCPIP^ox^ reduction) for up to 30 min. Temperature logging revealed that the sample temperature varied by <2 °C within the extended illumination periods. PSII-sample filled cuvettes in the spectrometer were continuously illuminated from the top side using a white-light lamp (Walz KL1500) with attenuation option, which was directed through a heat-protection filter (Schott KG5, ca. 400-700 nm transmission) to the cuvette by a ~20 cm light-guide (the full cuvette volume was homogenously illuminated). Several spectra (or time points) were recorded in the dark (prior to and after addition of, e.g., DCPIP), the light was switched on (or off) at indicated time points, and data were recorded on a PC linked to the spectrometer. Evaluation and fit analysis of absorption data was carried out using the Origin software (OriginLab). Light intensities at the sample center position were determined using a calibrated sensor device inserted in the spectrometer.

### X-ray absorption spectroscopy

XAS at the Mn K-edge was performed at beamline KMC-3 at the BESSY-II synchrotron (Helmholtz Zentrum Berlin) with the storage ring operated in top-up mode (250 mA), using a standard set-up as previously described^25,75^. A double-crystal Si[111] monochromator was used for energy scanning, the sample X-ray fluorescence was monitored with an energy-resolving 13-element germanium detector (Canberra), and samples were held in a liquid-helium cryostat (Oxford) at 20 K (in a 0.2 bar He heat-exchange gas atmosphere at an angle of 55 ° to the incident X-ray beam). The X-ray spot size on the sample was shaped by slits to about 1 (vertical) x 5 (horizontal) mm^2^, the X-ray flux was ~10^10^ photons s^-1^, the EXAFS scan duration was ~30 min. The energy axis was calibrated (±0.1 eV accuracy) using a Gaussian fit to the pre-edge peak (6543.3 eV) in the transmission spectrum of a permanganate (KMnO_4_) powder sample, which was measured in parallel to the PSII samples. For XAS data evaluation, up to 30 deadtime-corrected, energy-calibrated (I/I0) XAS monochromator scans (each on a fresh sample spot) were averaged and normalized XANES and EXAFS spectra were extracted after background subtraction as previously described using in-house software^76^. EXAFS simulations in *k*-space were carried out using inhouse software (SimX) and scattering phase functions calculated with FEFF9.077(S_0_^2^ = 0.8). Calculation of the filtered *R*-factor (*R*_F_, difference in %between fit curve and Fourier-backtransform of the experimental data in a 1-5 Å region of reduced distance)78facilitated evaluation of the EXAFS fit quality. Fourier-transforms of EXAFS spectra were calculated with cos windows extending over 10 % of both *k*-range ends.

### Sample preparation for XAS

Powder samples of manganese reference compounds (Mn oxides) were prepared from commercially available chemicals (MnCl_2_, Mn oxides) or from material (buserite, birnessite) that was kindly provided by the group of P. Kurz (Uni. Freiburg, Germany), diluted by grinding with boron-nitride (BN) to a level, which resulted in <15 % absorption at the K-edge maximum to avoid flattening effects in fluorescence-detected XAS spectra, loaded into Kapton-covered acrylic-glass holders, and frozen in liquid nitrogen. Aqueous MnCl_2_ (20 mM) samples were prepared at pH 7.0. Unless otherwise specified, PSII samples were prepared as follows: Mn-depleted PSII samples (3 mL) were prepared similar to the samples for optical absorption spectroscopy (see above), the pH was adjusted to the desired value, and samples were illuminated for 3 min at 1000 μE m^-2^ s^-1^ or kept in the dark as a control after addition of 240 μM MnCl_2_ and 60 μM PPBQ^ox^ (pH 7.5). Thereafter, the cuvette volume was rapidly mixed with ice-cold MES buffer (7 mL, pH 7.5, see above for ingredients) on ice in the dark, the pH was measured using a pH electrode and, if necessary, readjusted to the desired value (+/-0.1 pH units), the PSII membranes were pelleted by centrifugation (10 min, 20000 g, 2 °C), and kept on ice. Several of these sample types were rapidly merged on ice in the dark by loading (~30 μL) into XAS holders, which were immediately frozen in liquid nitrogen. Native PSII samples were prepared by pelleting of dark-adapted O_2_-evolving PSII membrane particles (~8 mg chlorophyll mL^-1^, pH 6.3), loading of the pellet material into XAS holders, and freezing in liquid nitrogen^73^. The shown XAS data for the electrodeposited Mn oxides has been collected in the context of earlier studies (16-18) and replotted.

## Data availability

All data needed to support the conclusions of this manuscript are included in the main text and SI Appendix.

## Author contributions

H.D. designed the study. P.C., S.F., J.H., N.O., and D.J.N. performed the experiments. P.C., M.H., and I.Z. analyzed the data. R.L.B., D.J.N., M.H., and H.D. wrote the manuscript.

## Acknowledgements

We thank I. Zizak, G. Schuck, and colleagues (Helmholtz Zentrum Berlin) for technical support in the X-ray experiments at the BESSY synchrotron. We thank P. Kurz (Universität Freiburg) for providing samples of birnessite and buserite. Financial support by the Deutsche Forschungsgemeinschaft (DFG) within SFB 1078 (project A4) and by the Einstein Foundation (Berlin, Einstein Fellow project awarded to R. Burnap) is gratefully acknowledged. Moreover, this study has been funded by the Deutsche Forschungsgemeinschaft (DFG, German Research Foundation) under Germany’s Excellence Strategy – EXC 2008/1 – 390540038 (Gefördert durch die Deutsche Forschungsgemeinschaft (DFG) im Rahmen der Exzellenzstrategie des Bundes und der Länder – EXC 2008/1 – 390540038).

