## Supplementary File for "Light-driven formation of high-valent manganese oxide by photosystem II supports evolutionary role in early bioenergetics"

**This PDF file includes:**

Supplementary text on experimental procedures
Figures S1 to S12
Tables S1 to S2
Supplementary text providing background information

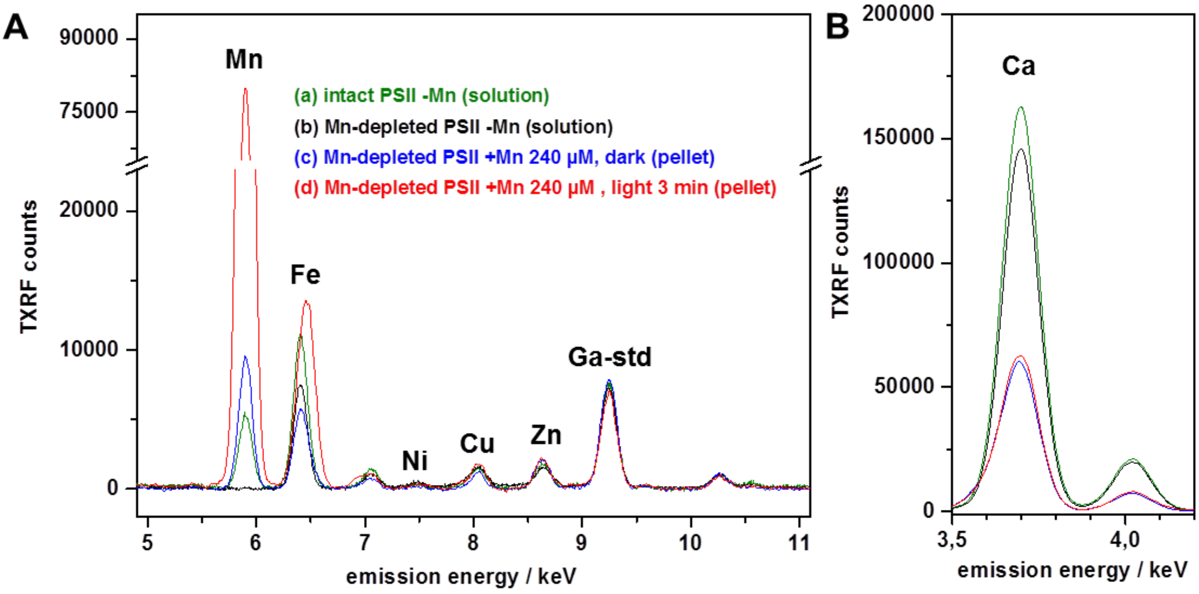


**Figure S1.** Metal content determination in PSII preparations by TXRF. A gallium concentration standard (Ga-std, 1 mg/L) was added to solutions of PSII membranes containing 1 mg/mL (~1 mM) chlorophyll (chl). Kα X-ray emission lines are labelled by element (respective unlabeled bands represent Kβ emission lines). +/-Mn denotes the presence or absence of additionally added manganese ($MnCl_2$). Respective Mn and Fe concentrations are given in Tables 1 and S1. **(A)** Spectra in the Mn to Ga region. **(a)** As-prepared PSII membrane particles, which were active in oxygen evolution, were pelleted and adjusted by buffer (pH 7) addition to a chl concentration of 1 mM. **(b)** Mn-depleted PSII membrane particles, which were inactive in oxygen evolution, were pelleted and adjusted by buffer (pH 7) addition to a chl concentration of 1 mM. **(c)** Mn-depleted PSII membrane particles (20 µg/mL / ~20 µM chl) were incubated for 3 min in darkness at 20 °C with 240 µM $MnCl_2$ in buffer (pH 8), pelleted, and adjusted by buffer addition to a chl concentration of 2 mM. **(d)** Mn-depleted PSII membrane particles (20 µg/mL / ~20 µM chl) were incubated for 3 min at 20 °C under continuous white light illumination (1000 µE $m^{-2}$ $s^{-1}$) with 240 µM $MnCl_2$ in buffer (pH 8), pelleted, and adjusted by buffer addition to a chl concentration of 2 mM. PSII membrane particles were pelleted by centrifugation at 50000 g for 12 min at 4 °C. **(B)** Spectra in the Ca region (spectra correspond to data in A). TXRF spectra were collected during ~10 min data acquisition (Bruker Picofox instrument) and metal contents were determined using the routines provided with the spectrometer. Note that the apparent shift of the Fe peak in (d) is due to the underlying large Kβ emission line of Mn. Spectra were normalized to the mean amplitude of the Ga standard Kα peak for comparison.

**Table S1:** Metal contents from TXRF in PSII preparations.[a]

| preparation | Mn [mg/L] [µM] | Fe [mg/L] [µM] | Ca [mg/L] [µM] | Mn/PSII | Fe/PSII | Ca/PSII |
|---|---|---|---|---|---|---|
| (a) intact PSII | 1.2±0.2<br>22±4 | 2.6±0.4<br>47±7 | 240.5±15<br>6000±380 | 4±1 | 9±2 | 1200±80 |
| (b) Mn-depleted PSII | 0.05±0.05<br><1±1 | 1.7±0.3<br>30±6 | 216.4±14<br>5400±350 | <0.2±0.2 | 6±2 | 1080±70 |
| (c) Mn-depl. PSII +240 µM Mn, dark | 2.0±0.4<br>36±7 | 1.4±0.4<br>25±7 | 88.2±6.0<br>2200±150 | 7±2 | 5±1 | 440±30 |
| (d) Mn-depl. PSII +240 µM Mn, light | 18±5<br>330±90 | 1.4±0.4<br>25±7 | 92.2±8.0<br>2300±200 | 65±19 | 5±1 | 460±40 |

[a]Data represent mean values of three repetitions (±standard deviation) of TXRF measurements as shown in Fig. S1. The concentration of the Ga standard was 1 mg/L (14.3 µM), the chlorophyll (chl) concentration was 1 mg/mL (~1 mM), Mn/PSII, Fe/PSII, and Ca/PSII values were calculated using the earlier determined value of about 200 chlorophylls per PSII reaction center [1,2]. Note that calcium concentrations and Ca/PSII values reflect predominantly the calcium in the used buffer (5 mM $CaCl_2$) and calcium bound unspecifically to the membrane proteins. In addition, concentration determination in comparison to the gallium standard is less accurate at the relatively low energy of the Ca emission lines. According to the XAS data (Fig. 5), PSII samples of type (c) contained about equal amounts of Mn(III) and Mn(IV) ions. Each PSII center therefore has transferred about 100 electrons, which would correspond to the double-reduction of about 50 DCPIP molecules per PSII. For ~0.1 µM PSII reaction centers (at ~20 µM chl) in the UV/vis assay of DCPIP (60 µM) reduction, the double-reduction of ~5 µM DCPIP, corresponding to a close to 10 % decrease of its absorption maximum at 604 nm thus was expected. This expectation is in reasonable agreement with the observed rapid decrease by ~0.1 units of the initial DCPIP absorption of ~0.8 (taking into account the chlorophyll background absorption) as shown in Fig. 3.

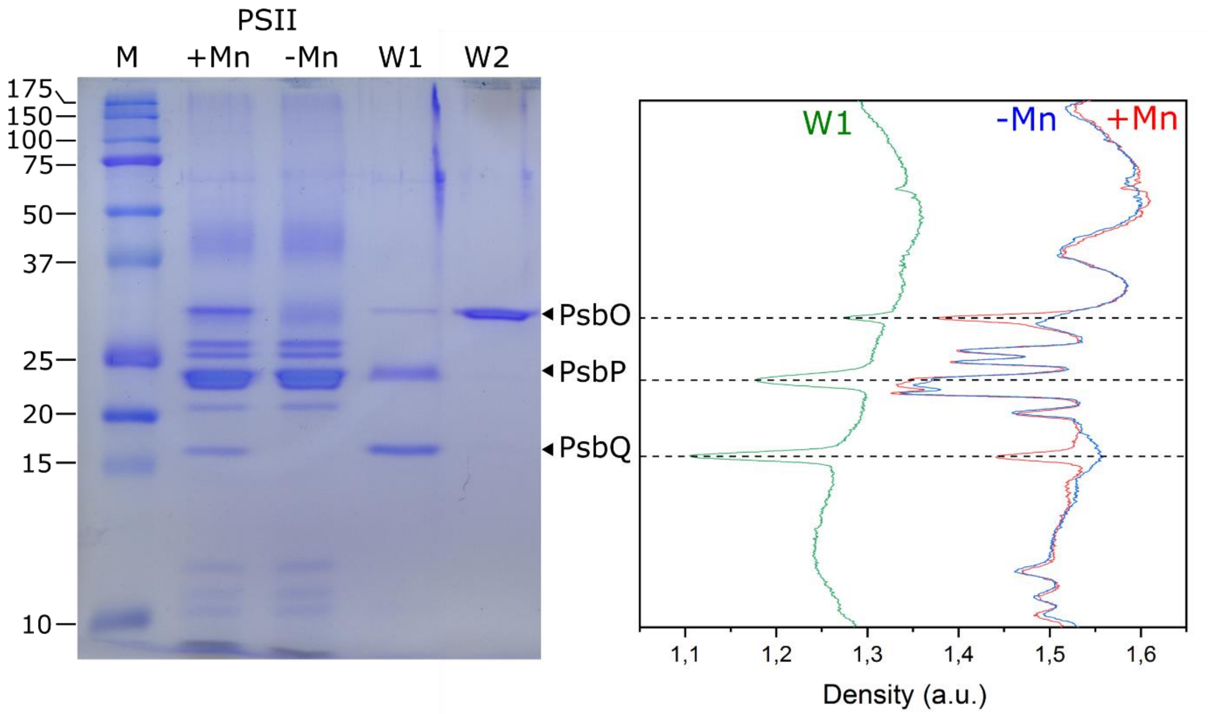


**Figure S2.** SDS-PAGE of PSII-enriched membrane particles and densitogram. The lanes represent: *M*, molecular weight marker; *+Mn*, PSII-enriched membrane particles directly after isolation; *-Mn*, Mn-depleted PSII-enriched membrane particles lacking the extrinsic proteins PsbO (33 kDa), PsbP (24 kDa), and PsbQ (18 kDa); *W1*, supernatant after washing with high-salt buffer containing TEMED; *W2*, proteins removed by washing with high-pH buffer. PSII samples with 1 µg chlorophyll and wash samples with 1 µg protein were loaded on the gel. The data show that the Mn-depletion and washing procedures had completely removed PsbQ and mostly PsbO and PsbP from the PSII protein complex (so that they appear in the supernatant after centrifugation) in samples used for the spectroscopic experiments. This is consistent with previous results by [3].

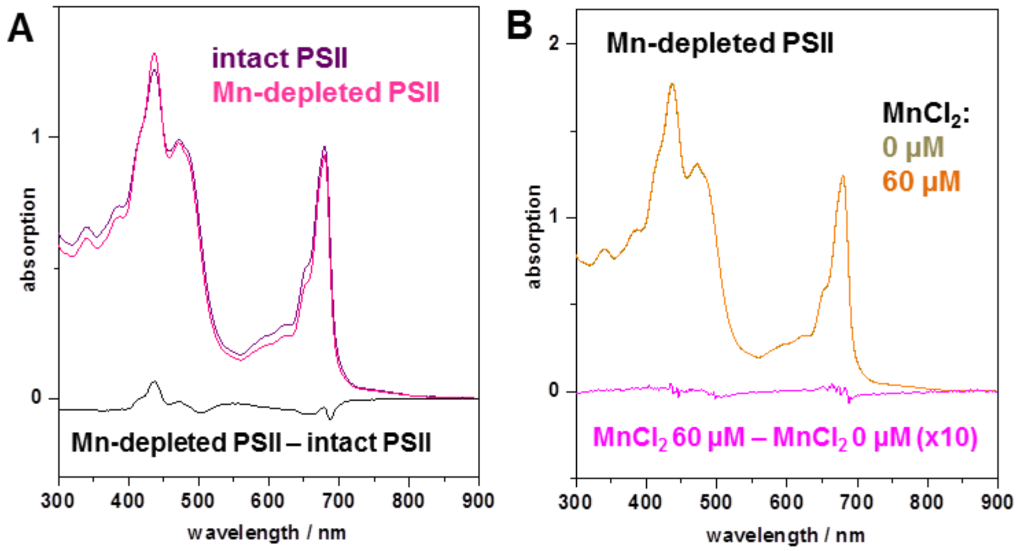


**Figure S3.** Optical absorption spectra of PSII preparations. **(A)** Comparison of intact and Mn-depleted PSII membrane particles and difference spectrum. Spectral differences in Mn-depleted PSII likely reflect partial removal of light harvesting complexes from the membranes during the manganese depletion procedure involving several washing (centrifugation) steps. **(B)** Mn-depleted PSII without (0 µM) and with 60 µM $MnCl_2$ and difference spectrum. Chlorophyll 20 µg/mL, pH 7.5 in (A) and (B). $MnCl_2$ has no significant absorption in the shown spectral region (300-900 nm).

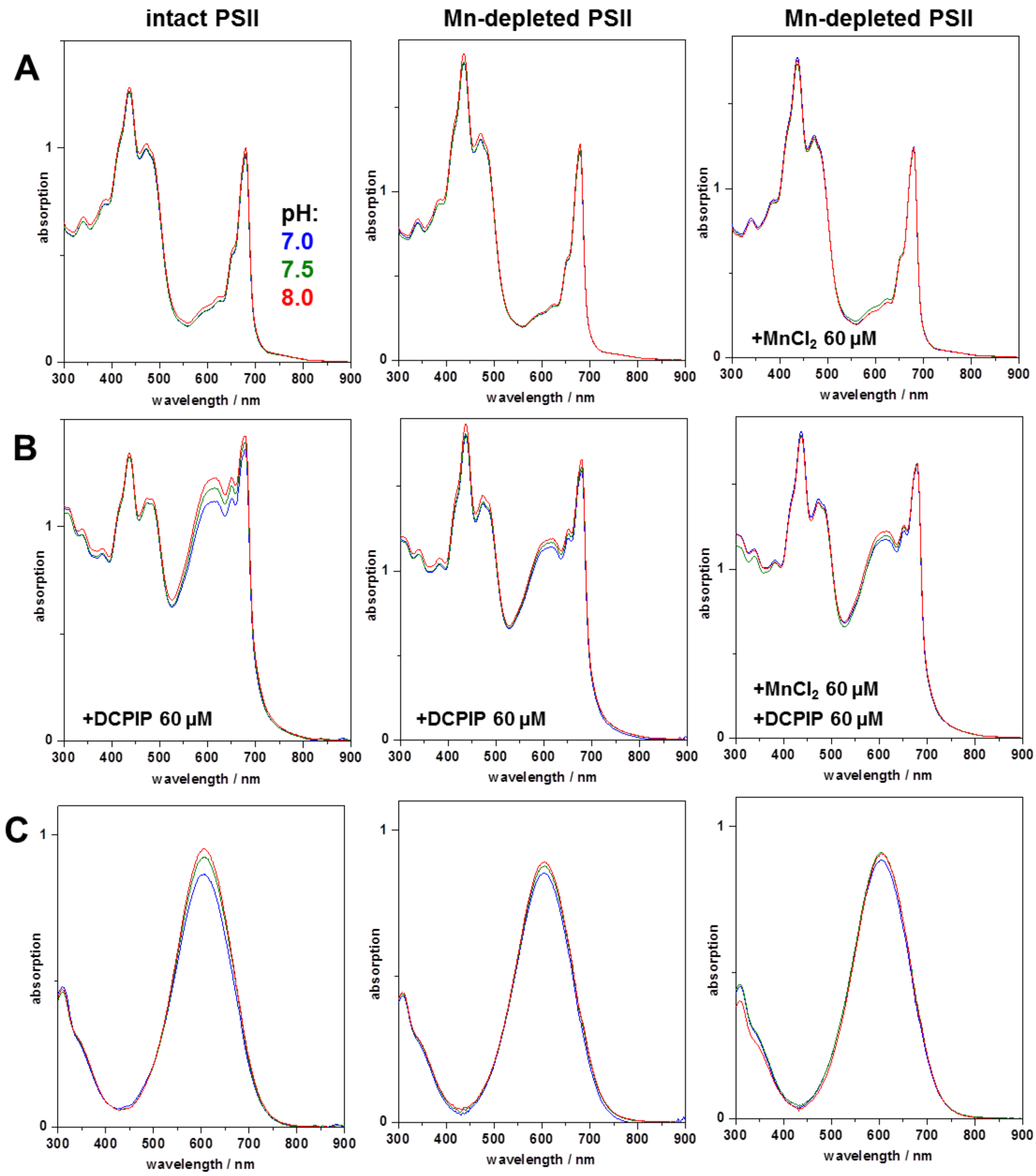


**Figure S4.** Absorption spectra of intact and Mn-depleted PSII. Spectra were collected on samples of PSII membrane particles without illumination (chlorophyll 20 µg/mL) at three pH values. **(A)** PSII preparations, **(B)** PSII preparations +DCPIP 60 µM, **(C)** (PSII +DCPIP) minus (PSII). Spectra in the right column were obtained for samples with addition of 60 µM $MnCl_2$. Note practically pH-independent spectra of the PSII preparations, no spectral contributions from $MnCl_2$, and an only slight pH-dependence (<5 % change) of the absorption maximum of oxidized DCPIP between pH 7.0 and 8.0 in (C).

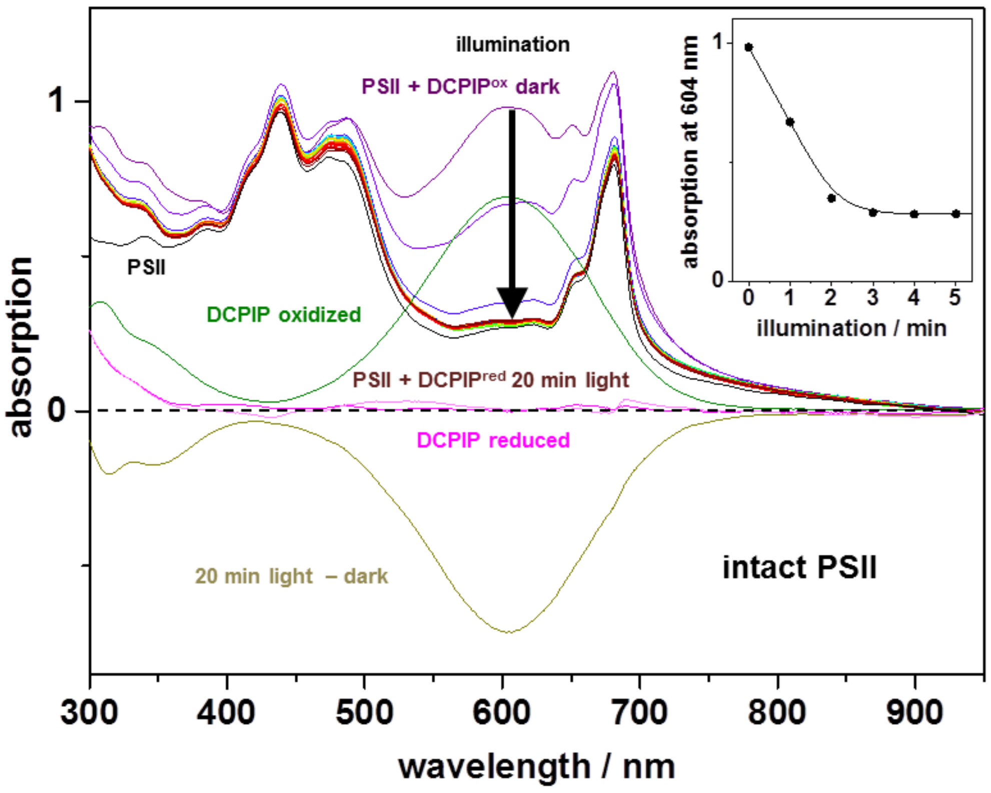


**Figure S5.** Absorption spectroscopy on DCPIP reduction by intact PSII. Oxygen-evolving PSII membrane particles were illuminated with saturating white light (1000 µE $m^{-2}$ $s^{-1}$, pH 7.0) and an absorption spectrum showing contributions from chlorophyll and DCPIP was recorded every 1 min. The dark spectra were obtained immediately prior to illumination. The spectrum of oxidized DCPIP (45 µM) was obtained before addition of PSII (20 µg $mL^{-1}$ chlorophyll), the difference spectrum (20 min light – dark) shows complete DCPIP reduction, and the spectrum of reduced DCPIP (magenta line) is the difference (20 min light – dark) minus (DCPIP oxidized) or the difference (light magenta line) of (20 min light) minus (PSII). The inset shows the absorption at the maximum of oxidized DCPIP (604 nm) as function of the illumination time.

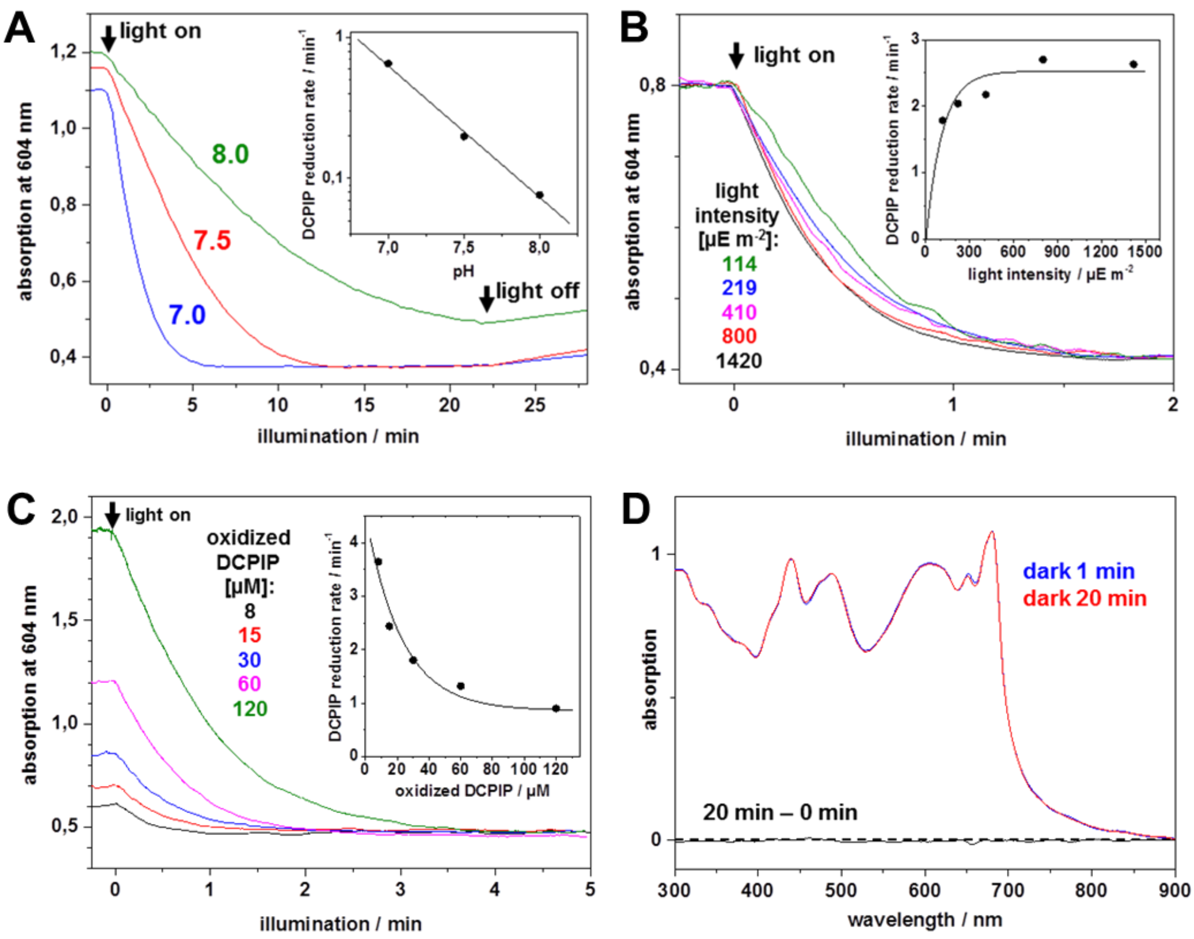


**Figure S6.** Kinetics of DCPIP reduction by intact PSII. DCPIP reduction was monitored at 604 nm (PSII membrane particles, 20 µg mL$^{-1}$ chlorophyll). **(A)** pH dependence. To oxygen-evolving PSII, 60 µM oxidized DCPIP were added in the dark and the light (1000 µE m$^{-2}$ s$^{-1}$) was switched on at t = 0 min and switched off at t = 22 min (arrows). DCPIP reduction was monitored in absorption spectra as in Fig. S5. The inset shows rate constants (circles, logarithmic scale) as determined from exponential fits of transients in the main panel together with a fit curve (line). **(B)** Light intensity dependence. To intact PSII, 30 µM oxidized DCPIP was added (pH 7). The inset shows rate constants of DCPIP reduction from single-exponential fits of data in the main panel together with a saturation curve (line). **(C)** DCPIP concentration dependence. To intact PSII, oxidized DCPIP was added (pH 7) and DCPIP reduction under illumination (~1000 µE m$^{-2}$ s$^{-1}$) was monitored. The inset shows rates of DCPIP reduction from exponential fits of data in the main panel together with an exponential decay curve (line). **(D)** Absence of reactions of Mn-depleted PSII in the dark. DCPIP (60 µM) and $MnCl_2$ (60 µM) were added to Mn-depleted PSII (pH 7) and spectra were collected after 1 min or 20 min darkness. No DCPIP reduction (or other changes) occurred in the dark.

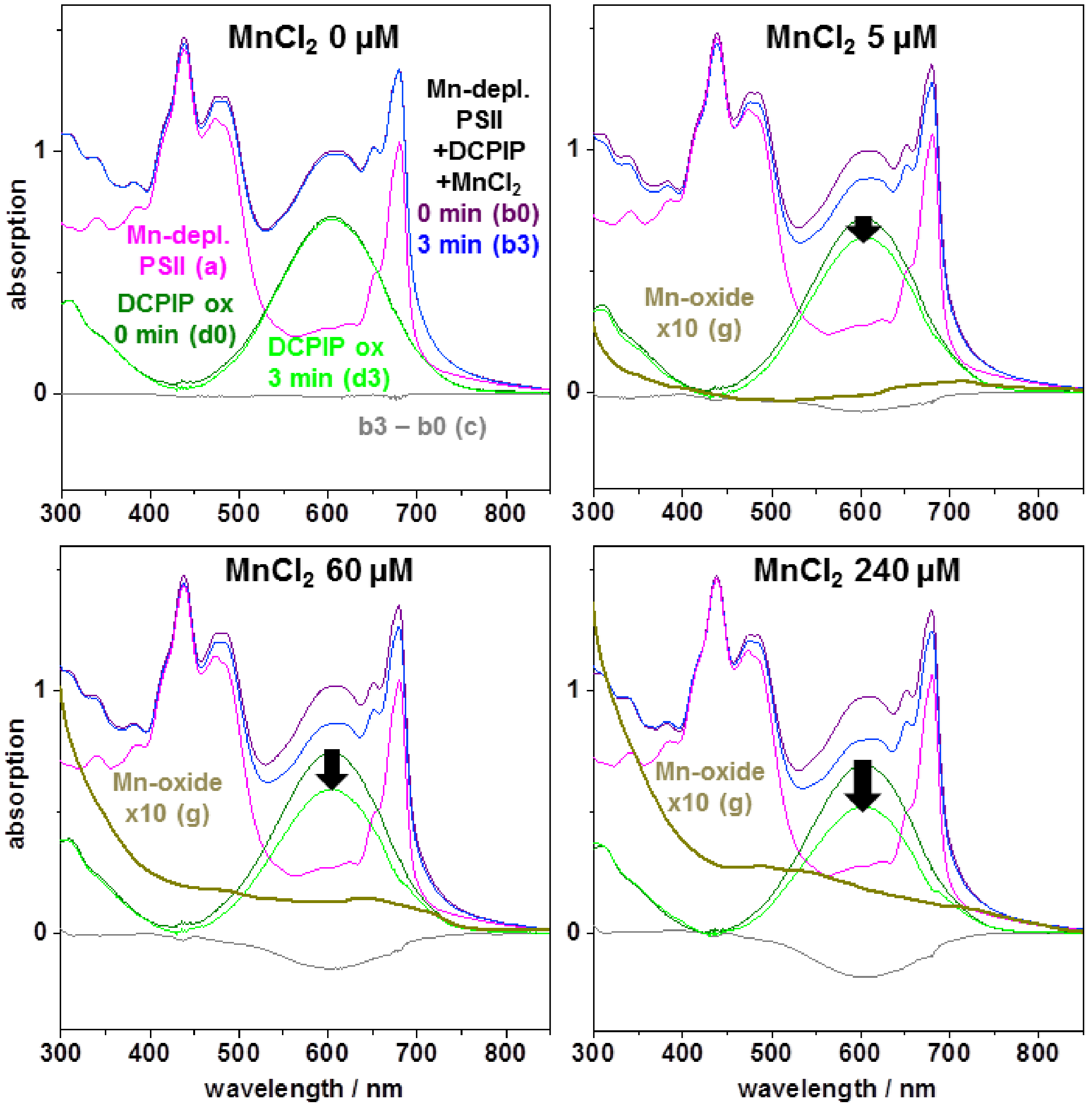


**Figure S7.** Light-driven electron transfer in Mn-depleted PSII for increasing $MnCl_2$ concentrations. Shown are optical absorption spectra of PSII (magenta spectra denoted a; 20 µg $mL^{-1}$ chlorophyll (chl), pH 7.0) to which 0-240 µM $MnCl_2$ and 60 µM $DCPIP^{ox}$ (dark-green spectra) were added (to yield spectra denoted b) and samples were illuminated with continuous white-light (1000 µE $m^{-2}$ $s^{-1}$) for 0 min or 3 min (purple and blue lines; 0 min, 3 min = spectra b0, b3; see also Fig. 3). The following difference spectra were calculated: c = b3 – b0 (difference after 3 min illumination); d0/3 = b0/3 – a (spectra of $DCPIP^{ox}$ after 0 min or 3 min illumination); e = c + 0.01a (0 µM $MnCl_2$; removal of chl bleaching contributions, leaving small remainders due to chl spectral changes); f = c + 0.01a + 0.10/0.20/0.27d (5/60/240 µM $MnCl_2$; chl bleaching removal and compensation for $DCPIP^{ox}$ decay); g = f – e (x10) (chl remainder removal yields spectra due to Mn oxide at 3 min illumination; smoothed by adjacent averaging of data points in a 50 nm range for clarity).

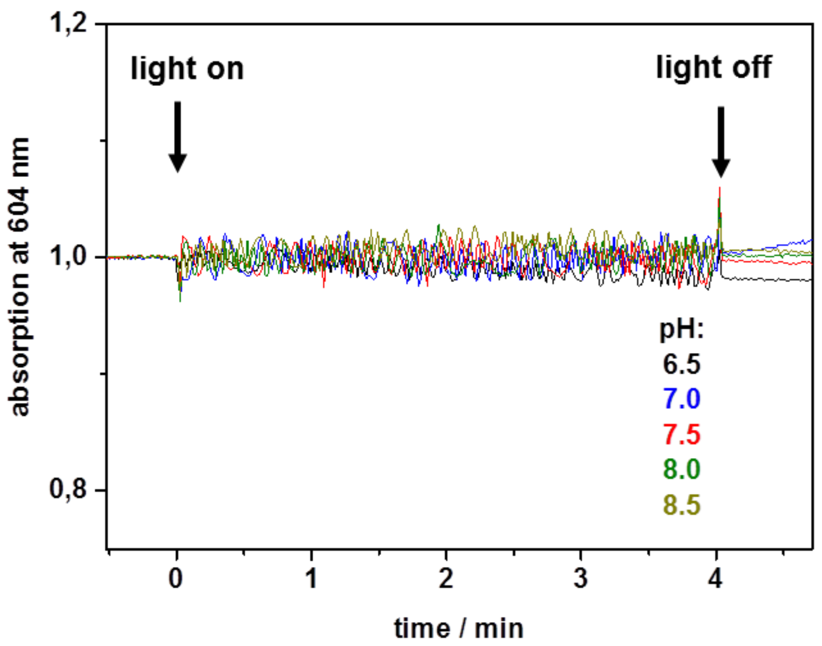


**Figure S8.** Absence of DCPIP reduction without PSII. Time traces of 604 nm absorption with 60 µM $DCPIP^{ox}$ and 240 µM $MnCl_2$ in buffers at the indicated pH values and in the absence of Mn-depleted PSII. Illumination (1000 µE $m^{-2}$ $s^{-1}$) of the solutions was started or stopped at the indicated time points (arrows). Note the absence of light-induced absorption changes, which indicate the absence of reactions of $MnCl_2$ with DCPIP in the absence of PSII.

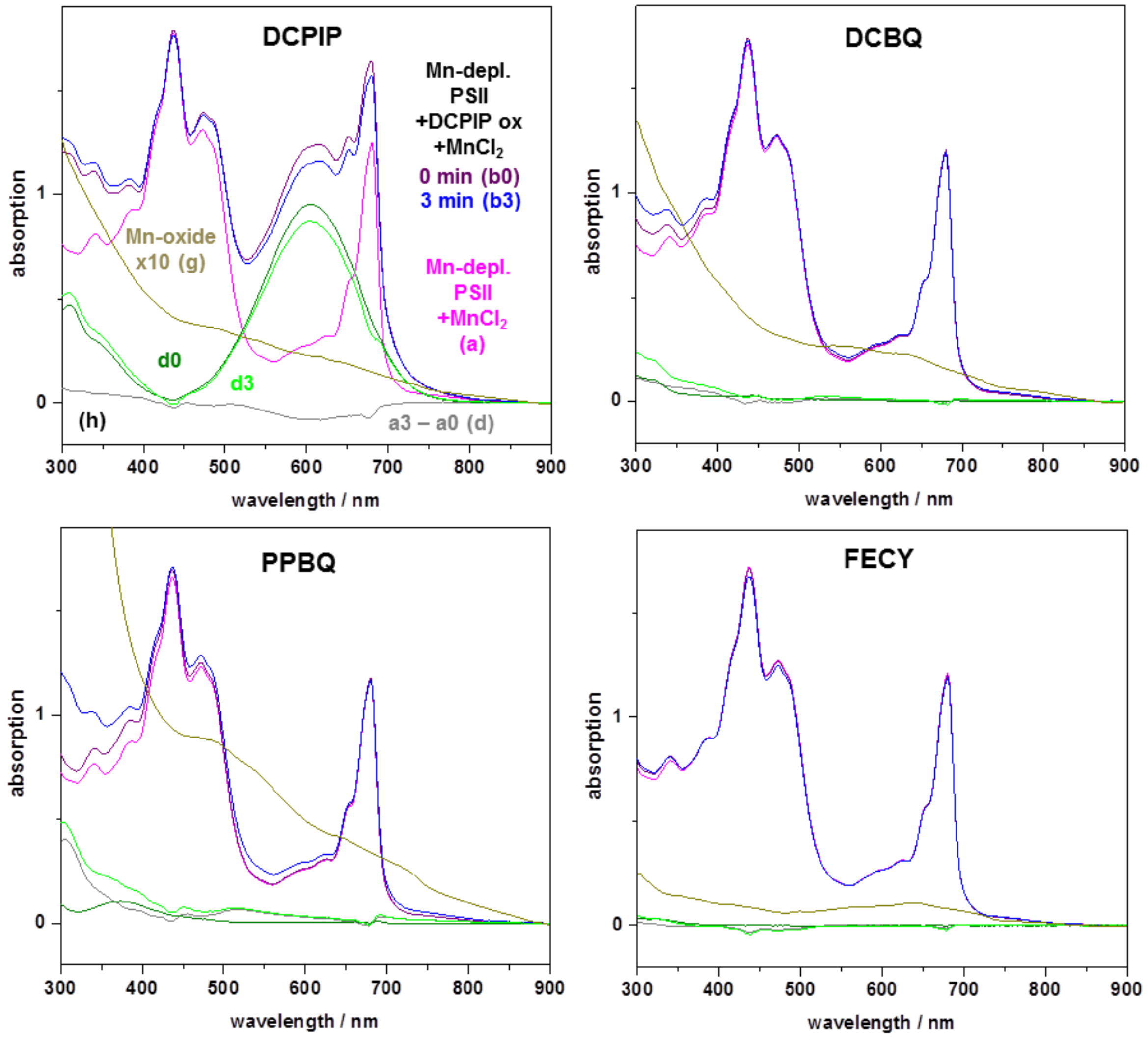


**Figure S9.** Formation of Mn oxide with different electron acceptors. Electron acceptors: DCPIP = 2,6-dichlorophenol-indophenol, DCBQ = 2,5-dichloro-1,4-benzoquinone, PPBQ = phenyl-p-benzoquinone, FECY = ferricyanide ($K_3Fe(III)(CN)_6$)). Optical absorption spectra of Mn-depleted PSII (20 µg $mL^{-1}$ chlorophyll, pH 8.0) to which 240 µM $MnCl_2$ (magenta spectra a) and initially oxidized electron acceptors (60 µM) were added and samples were illuminated with continuous white-light (4000 µE $m^{-2}$ $s^{-1}$) for 0 min (dark) or 3 min (0 min or 3 min illumination = spectra b0 (purple lines) and b3 (blue lines)). Difference spectra: c = b3 – b0 (spectral differences after 3 min illumination); d = b0 – a (initial oxidized acceptors); e = c + 0.01 a (0 µM $MnCl_2$; small spectral remainders due to chl spectral changes, see Fig. 3); f = c + 0.03/0.02/0.03/0.02 a + 0.09/0.10/0.28/0.05 d (DCPIP/DCBQ/PPBQ/FECY; approximate removal of chl bleaching and compensation for the decay of oxidized acceptors); g = f – e (x10) (removal of remainders results in dark-yellow spectra of reduced acceptors and oxidized manganese species after 3 min illumination; smoothed by adjacent averaging over 50 nm for clarity).

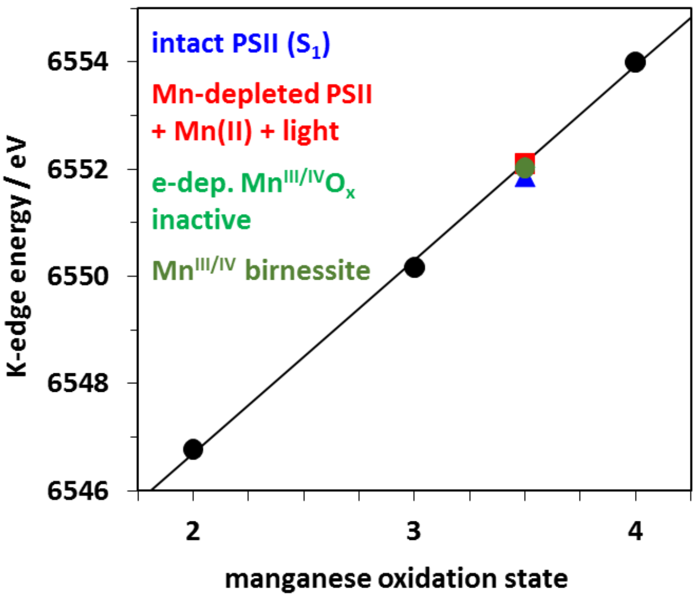


**Figure S10.** K-edge energies from XANES spectra vs. manganese oxidation state. Data points stem from K-edge spectra in Fig. 5A (e-dep. = electrodeposition) and were determined using the "integral method" for K-edge energy determination [4]. The line shows a linear fit ($E_{edge}$ = 6539.5 eV + 3.6 $Mn_{ox}$, $Mn_{ox}$ = Mn oxidation state) to the data points for the reference compounds ($Mn^{II}O$, α-$Mn^{III}_2O_3$, β-$Mn^{IV}O_2$). Note that K-edge energies of intact and Mn-depleted PSII correspond to a mean manganese redox level of +3.5, reflecting equal amounts of Mn(III) and Mn(IV) species.

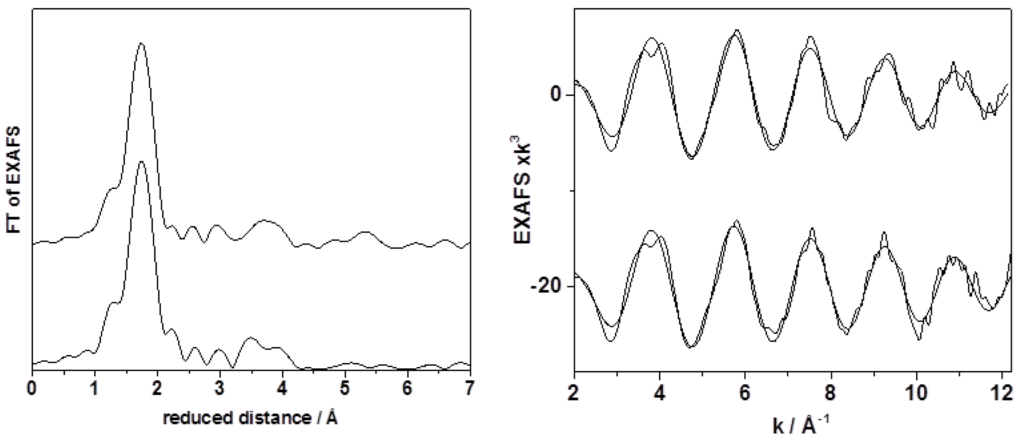


**Figure S11.** Mn K-edge X-ray absorption control experiment to explore spontaneous Mn oxide formation in the absence of PSII, at an eight-fold higher $Mn^{2+}$ concentration as used in the experiments with PSII (2 mM $MnCl_2$ dissolved in the aerobic standard buffer at pH 8 (top) and pH 7 (bottom); 60 µM PPBQ). The typical spectra of hexaquo $Mn^{2+}$ ions were obtained (6 Mn-O distances of ~2.15 Å); there are no indications for any spontaneous Mn oxide formation.

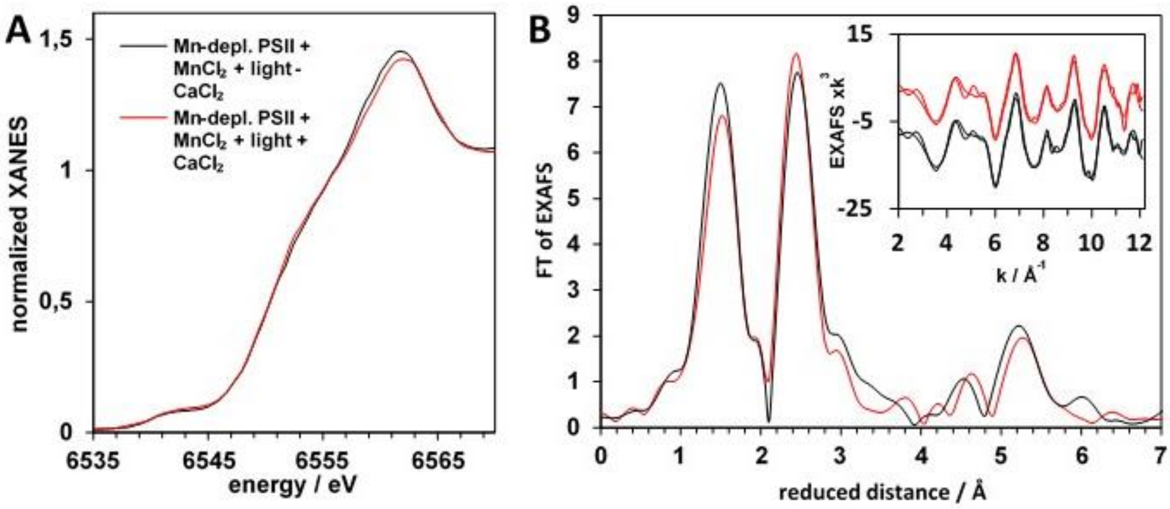


**Figure S12**. XAS spectra of PSII-bound Mn oxides formed in the presence and absence of $CaCl_2$ during light-induced Mn oxide formation. (A) Mn K-edge XANES. (B) Mn K-edge EXAFS. Main panel: Fourier transforms of spectra showing experimental data. Inset panel: EXAFS spectra in k-space (vertically shifted for clarity; thick lines, experimental data; thin lines, simulations with parameters detailed in Table S2). The illumination buffer contained 240 µM $MnCl_2$ and either 5 mM externally supplied $CaCl_2$ (red spectra) or no externally supplied $CaCl_2$, herein referred to as Ca-free buffer (black spectra). For the latter experiment, $CaCl_2$ had been removed from the suspension of Mn-depleted PSII by two rounds of centrifugation and resuspension steps in Ca-free buffer.

**Table S2:** EXAFS fit parameters.[a]

| shell | Mn-O | | | Mn-Mn | | | |
|---|---|---|---|---|---|---|---|
| sample | N [per Mn] | R [Å] | $2\sigma^2$ x$10^3$ [$Å^2$] | N [per Mn] | R [Å] | $2\sigma^2$ x$10^3$ [$Å^2$] | $R_F$ [%] |
| $Mn^{II}Cl_2$ | 6[§] | 2.19 | 10 | - | - | - | 36.0 |
| $Mn^{II}O$ | 6[§] | 2.19 (2.22) | 8 | 12[§] | 3.12 (3.14) | 6 | 27.4 |
| | | | | 6[§] | 4.50 (4.44) | | |
| | | | | 24[§] | 5.42 (5.44) | | |
| | | | | 12[§] | 6.39 (6.28) | | |
| α-$Mn^{III}_2O_3$ | 4.4[#] | 1.92 (1.96) | 10 | 6[§] | 3.10 (3.11) | 12 | 17.8 |
| | 1.6[#] | 2.22 (2.24) | | 6[§] | 3.56 (3.58) | 17 | |
| | | | | 6[§] | 4.70 (4.73) | 8 | |
| β-$Mn^{IV}O_2$ | 6[§] | 1.87 (1.89) | 7 | 2[§] | 2.85 (2.88) | 7 | 32.1 |
| | | | | 8[§] | 3.43 (3.43) | | |
| | | | | 5.1 | 5.04 (5.07) | | |
| birnessite $Mn^{III,IV}$ | 5.0[#] | 1.90 | 4 | 4.8 | 2.89 | 9 | 11.9 |
| | | | | 0.3 | 3.42 | | |
| | 1.0[#] | 2.30 | | 2.2 | 5.01 | | |
| | | | | 2.4 | 5.52 | | |
| electrodeposited inactive MnOx | 4.9[#] | 1.90 | 5 | 4.8 | 2.86 | 7 | 18.1 |
| | | | | 2.3 | 3.51 | | |
| | 1.1[#] | 2.33 | | 2.5 | 5.02 | | |
| | | | | 6.0 | 5.55 | | |
| electrodeposited active MnOx (MnCat) | 5.1[#] | 1.89 | 8 | 2.3 | 2.87 | 9 | 15.0 |
| | | | | 1.6 | 3.45 | | |
| | 0.9[#] | 2.30 | | 1.4 | 5.01 | | |
| | | | | 1.7 | 5.48 | | |
| intact PSII ($S_1$) | 3.6[#] | 1.83 | 12 | 1.5 | 2.73 | 4 | 16.8 |
| | 2.4[#] | 2.01 | | 0.6 | 3.25 | | |

Mn-depleted PSII (+3 min light, +240 µM $MnCl_2$, 5 mM $CaCl_2$, 60 µM PPBQ)

| | | | | | | | |
|---|---|---|---|---|---|---|---|
| (a) | 4.7# | 1.90 | 6 | 3.4 | 2.87 | 6 | 21.5 |
| | | | | 0.7 | 3.46 | | |
| | 1.3# | 2.25 | | 1.9 | 5.01 | | |
| | | | | 4.9 | 5.53 | | |
| (b) | 4.6# | 1.91 | 5 | 2.7 | 2.87 | 6 | 25.6 |
| | | | | 0.8 | 3.45 | | |
| | 1.4# | 2.22 | | 1.2 | 4.97 | | |
| | | | | 4.2 | 5.52 | | |
| (c) | 5.2# | 1.90 | 6 | 4.1 | 2.89 | 5 | 12.8 |
| | | | | 0.8 | 3.54 | | |
| | 0.8# | 2.28 | | 1.9 | 4.99 | | |
| | | | | 4.6 | 5.57 | | |
| mean | 5.0# | 1.90 | 6 | 3.5 | 2.87 | 6 | 13.5 |
| | | | | 0.7 | 3.49 | | |
| | 1.0# | 2.25 | | 1.5 | 4.99 | | |
| | | | | 4.2 | 5.54 | | |

Mn-depleted PSII (+3 min light, +240 µM $MnCl_2$, with/without $CaCl_2$, PPBQ)

| shell | Mn-O | | | Mn-Mn | | | |
|---|---|---|---|---|---|---|---|
| sample | N [per Mn] | R [Å] | $2\sigma^2$ x$10^3$ [Å$^2$] | N [per Mn] | R [Å] | $2\sigma^2$ x$10^3$ [Å$^2$] | $R_F$ [%] |
| Mn-depleted PSII (+1 min light, +240 µM $MnCl_2$, +30 µM PPBQ + 5 mM $CaCl_2$) | 4.8# | 1.89 | 8 | 3.5 | 2.86 | 4 | 18.4 |
| | | | | 0.6 | 3.52 | | |
| | 1.2# | 2.24 | | 1.7 | 5.01 | | |
| | | | | 4.6 | 5.56 | | |
| Mn-depleted PSII (+1 min light, +240 µM $MnCl_2$, +30 µM PPBQ - $CaCl_2$) | 4.9# | 1.89 | 7 | 3.5 | 2.87 | 5 | 12.0 |
| | | | | 1.1 | 3.52 | | |
| | 1.1# | 2.28 | | 1.4 | 4.98 | | |
| | | | | 4.2 | 5.56 | | |

[a]*N*, coordination number; *R*, interatomic distance; $2\sigma^2$, Debye-Waller parameter; $R_F$, fit error sum (for reduced distances of 1-5 Å). Fit restraints: [§]fixed parameter, [#]*N*-values coupled to yield a sum of 6. [$]These distances in intact PSII contain contributions from Mn-Ca vectors in the $Mn_4CaO_5$ complex. Parameters correspond to spectra in Figs. 5 and S11. $MnCl_2$ was measured as 10 mM aqueous solution. For MnO, α-$Mn_2O_3$ and β-$MnO_2$ spectra commercial compounds were used and mixed with BN in ~1:20 ratio (supplier and purity on trace metal basis: $Mn_2O_3$ Sigma-Aldrich, 99.9%; $MnO_2$, Carl Roth, 99.995%; MnO, Sigma-Aldrich, 99%).

The fit parameters for the Mn oxides comply with earlier EXAFS results on the respective and related species [5-10]. In parentheses, (average) distances determined by crystallography are given for $Mn^{II}O$ [11] (Crystallography Open Database ID 1514105), α-$Mn^{III}_2O_3$ [12] (Crystallography

Open Database ID 1514103), and β-$Mn^{IV}O_2$ [13] (Crystallography Open Database ID 1514117). Good agreement between crystallographic and EXAFS distances verifies the intactness of the commercial Mn oxides. The birnessite spectrum corresponds to the $K_{0.31}$-birnessite spectrum published in [9], where further characterization of the material is provided. The spectra of electrodeposited active and inactive MnOx correspond to the spectra measured under open circuit conditions and published previously in [7].

**Additional text:**

**Background on $Mn^{2+}$ oxidation by PSII and hypotheses on the evolution of the oxygen-evolving complex (OEC) of PSII**

**$Mn^{2+}$ oxidation by PSII:** The characteristics of electron donation to PSII by $Mn^{2+}$ are very well studied, but to our knowledge, an analysis of the structural and chemical configuration of the Mn products of the reaction, except in the case of photoactivation and the reformation of the native water-oxidizing Mn cluster, has not been described until now. Early studies on the properties of electron transport in the PSII reaction center showed that $Mn^{2+}$ served as an electron donor to Mn-depleted PSII [14-18]. Of particular relevance, several studies showed that added $Mn^{2+}$ could support sustained DCPIP photoreduction in Mn-depleted chloroplast and PSII preparations [18]. Hoganson and Babcock later showed $Mn^{2+}$ was directly oxidized by the redox active tyrosine, $Y_Z$, of the D1 polypeptide [19] providing the basis for the current structural understanding of the pathway. Kinetic analysis of the competition between $Mn^{2+}$ and other donors has subsequently provided further information on the affinity characteristics of $Mn^{2+}$ binding [15,20,21]. While advancing the understanding of the binding of Mn ions to PSII, these various studies did not, however, provide information on the physiochemical nature of the products of the $Mn^{2+}$ photochemical oxidation reaction as reported here. These studies did, however, lead to the discovery of photoactivation, which is the photochemical synthesis of the catalytically active $Mn_4CaO_5$ [22-26]. Unlike the very large arrays of Mn oxides described in the present work, photoactivation results in the formation of only the $Mn_4CaO_5$. Although much remains to be learned about photoactivation, important information on the nature of the structural intermediates has been published including EPR characterizations of early assembly intermediates [27,28].

The pioneering studies of George Cheniae showed that when photoactivation is performed under sub-optimal conditions, such as in the absence of $Ca^{2+}$, inactive Mn is accumulated. Under these conditions, the yield of Mn clusters that were capable of water-splitting decreased, yet Mn was found to be tightly bond in an EDTA unextractable, EPR silent form [29]. Cheniae referred to this form of photochemically assembled, but catalytically inactive Mn as "inappropriately bound ligation of $Mn^{\geq 3+}$" [29] and the formation of these deposits corresponded to a type of photoinactivation preventing proper cluster assembly. However, these scientists did not characterize the form of the metal beyond estimating the number of atoms per reaction center nor did they speculate on whether or not these corresponded to Mn oxides. In retrospect, it appears likely that they were indeed making the Mn oxides in their reactions, although their comparatively small size would have made the structural characterization more difficult than what is now observed. It is worth noting that the Mn oxide structures considered by Russell and Hall [30] as well as by Sauer and Yachandra [31] were argued as abiotically formed and subsequently incorporated into a reaction center that was a precursor to modern PSII, which contrasts with the alternative proposal that the formation of the oxides occurs via photooxidation of $Mn^{2+}$ at a reaction center.

**Hypotheses on the evolution of the OEC:** The evolutionary transition from anoxygenic photosynthesis to oxygenic photosynthesis would have involved the acquisition of metal binding features on the donor side of primitive anoxygenic photosynthetic reaction centers.

Allen and co-workers have tested the hypothesis that it is possible to modify the redox and metal binding properties of anoxygenic purple bacterial reaction centers to produce modified complexes capable of oxidizing $Mn^{2+}$ [32] Blankenship and Hartman proposed that the ancestral proto-PSII interacted with a Mn catalase [33]. Another class of proposals is that the $Mn_4CaO_5$ originates by the incorporation of a preformed mineral for the origin of a ‘preformed’ abiotic core cluster. Dismukes, Klimov and colleagues proposed that reaction center proteins acquired an $O_2$-evolving Mn cluster by binding of a tetramanganese-bicarbonate cluster derived from the Mn-bicarbonate clusters present in the environment [34]. Russell and Hall performed seminal comparative analyses of the $O_2$-evolving $Mn_4CaO_5$ with naturally occurring inorganic Mn oxides [30]. They suggested photochemically formed $MnO_2$ precipitates in the early ocean were recruited and later modified to the $O_2$-evolving Mn cluster. They reasoned that the $O_2$-evolving Mn cluster may have derived from a coating of manganese precipitates containing $Mn^{4+}$ that may have served as an electron acceptor and/or to protect from hard ultraviolet damage and "Thus, a cluster of ranciéite may have contributed the “ready-made” [$CaMn_4$] structure that was co-opted by one of the reaction centers, though if so the Mn-Mn distances of 2.9Å characteristic of the cluster constrained in ranciéite must have been modified to a conformation more typical of hollandite where two Mn-Mn distances are 2.7Å and two are 3.3Å" [30]. Sauer and Yachandra postulated that MnO was incorporated and gave a ‘catalase’ activity, releasing hydrogen peroxide [31]. Later, the pre-PSII complex was modified to utilize and assemble water-splitting Mn clusters: "The evolution of the modern complex, however, presumably began in the absence of the refined PS II protein binding site and fully developed mechanism for photo-oxidizing $Mn^{2+}$ during its incorporation. It is well known that solid $MnO_2$ exhibits pronounced ‘‘catalase’’ activity in its ability to increase the rate of decomposition of $H_2O_2$ into $H_2O$ and $O_2$ by many orders of magnitude." This falls into a class of hypotheses that conjecture that Mn oxides available in the environment were recruited into the ancient photochemical reaction center and subsequently modified.

Another class of proposals for the evolutionary origin of the $O_2$-evolving $Mn_4CaO_5$ hypothesizes that the metal cluster is a modification of metabolic adaptation of ancient microbial photosynthetic bacteria that utilized metals, first $Fe^{3+}$ and later $Mn^{2+}$, as a source of metabolic reductant. Excellent recent synopsis on the current hypotheses regarding the evolution of photosynthetic reaction centers from a physiological perspective can be found in [35]. Zubay [36] and subsequently, Fischer et al. [37] proposed that an early precursor of PSII utilized solvated $Mn^{2+}$ as a source of electrons for metabolism, hypothesizing that the resultant by-products were Mn oxides. These were hypothesized to be an intermediate to the evolution to a modified form of the Mn oxide that was catalytic and had become capable of supplying electrons from a secondary donor such as water. Fischer went on to assign this process to the origin of certain geologic Mn deposits, but his studies were focused more on the geologic processes and did not obtain evidence for hypothesized Mn oxide formation by photochemical reaction centers [37]. Thus, the formation of oxides by PSII, though hypothesized had not been shown before and that is the experimental novelty we now report.
